# Individual bitter-sensing neurons in *Drosophila* exhibit both ON and OFF responses that influence synaptic plasticity

**DOI:** 10.1101/2020.08.25.266619

**Authors:** Anita V. Devineni, Julia U. Deere, Bei Sun, Richard Axel

**Author notes:** Correspondence (A.V.D.) or (R.A.).

## Abstract

The brain creates internal representations that translate sensory stimuli into appropriate behavior. Most studies of sensory processing focus on which subsets of neurons are activated by a stimulus, but the temporal features of the neural response are also important for behavior. In the taste system, the timing of peripheral sensory responses has rarely been examined. We investigated the temporal properties of taste responses in *Drosophila melanogaster* and discovered that different types of taste sensory neurons show striking differences in their response dynamics. Strong responses to stimulus onset (ON responses) and offset (OFF responses) were observed in bitter-sensing neurons in the labellum, whereas bitter neurons in the leg and other classes of labellar taste neurons showed only an ON response. Individual bitter labellar neurons generate both the ON and OFF responses through a cell-intrinsic mechanism that requires canonical bitter receptors. The bitter ON and OFF responses at the periphery are propagated to dopaminergic neurons that innervate the mushroom body and mediate aversive learning. When bitter is used as a reinforcement cue, the bitter ON and OFF responses can drive opposing types of synaptic plasticity and the effect of the OFF response dominates, likely due to the rapid and preferential habituation of the ON response. Together, these studies characterize novel features of neural responses in the taste system and reveal their importance for neural circuit function.

## INTRODUCTION

Organisms have evolved diverse mechanisms to recognize sensory information in the environment and transmit this information to the brain, where it is processed to create an internal representation of the external world. This internal representation must translate stimulus features into appropriate behavioral output. The efficient translation of external stimuli into behavior engages sensory neurons that may respond to the same stimulus in different ways. For example, ON cells respond to the onset of a stimulus or an increase in its intensity, whereas OFF cells respond to stimulus offset or decreased intensity. In most sensory systems ON cells predominate. OFF responses have been observed in visual, auditory, somatosensory, and chemosensory pathways (Abraira and Ginty, 2013; Hubel and Wiesel, 1961; Luo and Katz, 2001; Sato, 1976; Xu et al., 2014). ON-OFF cells that respond to both stimulus onset and offset are also observed (Nagel and Wilson, 2016; Pei et al., 2009; Qin et al., 2007; Scholl et al., 2010).

The disappearance of a stimulus may be as relevant to an organism as its appearance, and animals must respond to both events robustly. An OFF response enables the nervous system to detect and interpret stimulus decrements more rapidly and reliably than the decay of the ON response. Computational modeling suggests that ON-OFF systems provide more efficient information coding at low firing rates and under conditions of higher noise (Gjorgjieva et al., 2014). Thus OFF responses have been proposed to underlie the perception of stimulus duration, disappearance, and gaps in a stream of stimuli (Bregman et al., 1994; Liang et al., 2008; Mazor and Laurent, 2005; Xu et al., 2014).

OFF responses may also elicit behaviors distinct from ON responses. For example, the onset of an attractive odor causes an animal to orient upwind, whereas odor offset induces local search behaviors such as turning (Alvarez-Salvado et al., 2018; Kennedy and Marsh, 1974; van Breugel and Dickinson, 2014). In *C. elegans*, for instance, the AWA olfactory neuron is activated at odor onset and suppresses turning to promote forward locomotion, while the AWC olfactory neuron is activated only at odor offset and promotes turning (Chalasani et al., 2007; Larsch et al., 2015). The ability of ON and OFF responses to elicit different behaviors at stimulus onset and offset is most easily accomplished if they occur in different populations of cells. Many systems show this segregation of ON and OFF responses into different cells, including the visual system (Hubel and Wiesel, 1961; Westheimer, 2007) and the *C. elegans* olfactory system (Chalasani et al., 2007; Larsch et al., 2015). In other cases individual cells show both ON and OFF responses (Scholl et al., 2010; Xu et al., 2014), requiring more complex processing to interpret these two responses differently and elicit different behaviors.

In the taste system, different taste modalities are usually encoded by separate populations of sensory cells and drive innate behavioral responses (Liman et al., 2014). For example, sugar-sensing cells drive appetitive feeding responses, whereas bitter-sensing cells elicit aversion and suppress feeding. Insects have evolved a complex peripheral taste system and possess taste sensory neurons in multiple organs including the legs, wings, and labellum (the distal segment of the proboscis) (Scott, 2018). These external taste neurons enable insects to rapidly sample different substrates as they navigate an environment, and insects use taste cues to guide ongoing behaviors including feeding as well as locomotion, egg-laying, grooming, and courtship (Corfas et al., 2019; He et al., 2019; Joseph and Heberlein, 2012; Thistle et al., 2012; Thoma et al., 2016; Yanagawa et al., 2014). Thus the presence of ON and OFF responses to tastants may be important for the generation of behavior in insects. A study in moths reported that second-order taste neurons show dynamic, time-varying responses, including OFF responses (Reiter et al., 2015). However, the timing of gustatory responses has not been examined in the fruit fly *Drosophila melanogaster*.

In this study we investigated the timing of neuronal responses in the *Drosophila* taste system. We found that different taste sensory neurons show striking differences in their response dynamics. OFF responses were observed in bitter-sensing neurons of the labellum, but not in bitter neurons of the leg or in other labellar taste neurons that detect different taste modalities. The bitter OFF response depends on stimulus identity and concentration and is more sensitive to these features than the ON response. Individual labellar bitter neurons show both ON and OFF responses, and genetic and neuronal manipulations suggest that the OFF response is generated cell-intrinsically through canonical bitter receptors. These organ-specific response dynamics are propagated to downstream dopaminergic neurons that innervate the mushroom body and mediate aversive learning. We show that the bitter OFF response in these neurons drives synaptic plasticity when bitter is used as a reinforcement cue. Together, these studies identify ON-OFF cells in the fly taste system and reveal the importance of these dynamics on neural circuit function.

## RESULTS

### Taste neurons vary in the temporal properties of their responses

We analyzed the timing of *Drosophila* taste responses by performing calcium imaging at the axon terminals of sensory neurons, located within the subesophageal zone (SEZ) of the brain (Figure 1A). Small drops of tastants were delivered to the labellum or foreleg using a custom-built taste delivery system. Labellar bitter-sensing neurons responded strongly to bitter stimuli such as quinine, denatonium, and lobeline (Figure 1A-C). These bitter neurons showed strong peaks in calcium activity when the stimulus was presented (an ON response) and removed (an OFF response). The magnitude of the OFF response was often similar or larger than that of the ON response. For example, quinine elicited a peak ON response of ∼60% ΔF/F and a peak OFF response of ∼90% ΔF/F. The dynamics of the ON and OFF responses were remarkably consistent across trials, and both the ON and OFF responses were closely time-locked to the stimulus (Figure 1B). When we varied the stimulus duration from 3 to 10 sec, the OFF response shifted in time to correspond with the time of bitter offset (Figure 1C). The magnitude of the OFF response was generally similar for different stimulus durations (Figure 1C). These results demonstrate that labellar bitter-sensing neurons respond to bitter with both an ON and an OFF response.

**Figure 1:**
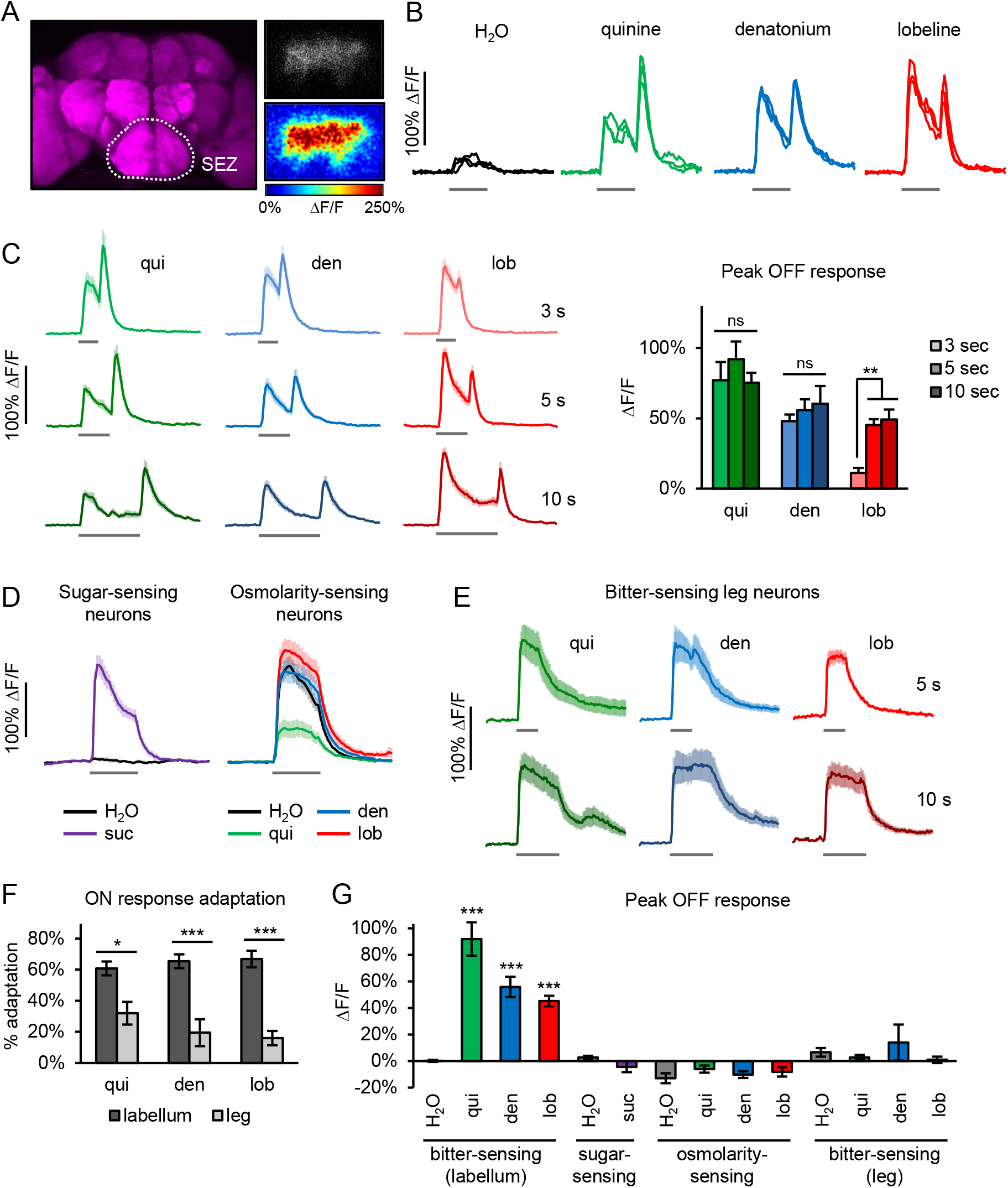
The timing of taste sensory responses varies by modality and organ. (A) Left: location of the SEZ in the brain, where GCaMP imaging of taste sensory axons was performed. Top right: image of bitter sensory axons expressing GCaMP. Bottom right: heatmap showing example GCaMP response to denatonium stimulation (average over 3 trials). (B) Labellar bitter-sensing neurons showed ON and OFF responses to 5 sec bitter stimulation. Graphs show GCaMP traces for individual trials from one fly (same example shown in panel A). (C) Labellar bitter-sensing neurons stimulated with bitter tastants for 3, 5, or 10 sec showed ON and OFF responses (n ≥ 8 trials, 3 flies). Left: average GCaMP traces. Right: peak OFF response for each condition. (D) Labellar sugar-sensing neurons (n = 9 trials, 3 flies) and osmolarity-sensing neurons (n = 18 trials, 6 flies) showed ON responses but no OFF response (5 sec stimulation). (E) Bitter-sensing neurons in the leg showed ON responses but no OFF response (n ≥ 9 trials, 3-5 flies). Non-responsive trials were excluded from these averages. (F) Labellar bitter neurons showed greater adaptation of the ON response than tarsal bitter neurons (same data as panels C and E for 5 sec stimulation). (G) Comparison of peak OFF responses across different types of taste neurons using 5 sec stimulation. Only labellar bitter neurons showed significant OFF responses. *Gal4* lines used to express *UAS-GCaMP6f* were *Gr98d-Gal4* (bitter-sensing neurons), *Gr64f-Gal4* (sugar-sensing neurons), or *ppk28-Gal4* (osmolarity-sensing neurons). *p<0.05, **p<0.01, ***p<0.001, one-way ANOVA followed by Tukey’s post-test (panel C), t-test comparing groups (panel F), or one-sample t-test vs. 0 (panel G; only significant values greater than zero are noted). See also Figure S1. In all figures, grey bars denote stimulus presentation. Unless otherwise specified, all graphs in this paper represent mean ± SEM.

We next examined whether other classes of labellar taste neurons also show OFF responses. Sugar-sensing neurons displayed a strong ON response but no OFF response when stimulated with sucrose (Figure 1D, 1G). We also imaged the osmolarity-sensing neurons, which respond to water as well as other low osmolarity solutions (Cameron et al., 2010), including the same bitter solutions that were used to activate bitter-sensing neurons. The osmolarity-sensing neurons showed strong ON responses but no OFF responses (Figure 1D, 1G). These results indicate that the OFF response is specific to the bitter taste modality.

We then imaged the responses of bitter neurons in the leg (tarsus) (Ling et al., 2014), which project to a different region of the SEZ than labellar bitter neurons (Wang et al., 2004). Bitter neurons in the leg showed strong ON responses, but no OFF response, to all three bitter compounds tested (Figure 1E, 1G, and S1A). However, we did observe an OFF response to one of the three compounds in just one of the 9 flies whose tarsal bitter neurons were imaged (Figure S1B-C). We also observed that the bitter ON response in labellar neurons, but not tarsal neurons, showed a strong decay during the stimulus presentation, termed adaptation (Figure 1C, 1E-F, and S1A). In labellar bitter neurons, the response to a 5 sec bitter stimulus decayed by at least 60% by the end of the stimulation, whereas the response in tarsal neurons decayed by only ∼15-30% (Figure 1C, 1E-F, and S1). With 10 sec bitter stimulation the labellar bitter response often decayed to near baseline (Figure 1C). The dynamics of these responses suggest that bitter neurons in the leg and labellum encode the same taste stimuli in fundamentally different ways: the leg signals the presence or absence of bitter, whereas the labellum signals bitter onset and offset.

### The bitter OFF response is present in all classes of labellar bitter neurons and depends on stimulus identity and concentration

Four classes of labellar bitter neurons have been identified based on the combination of bitter receptors that they express: S-a, S-b, I-a, and I-b (Weiss et al., 2011). Our initial experiments (Figure 1) focused on just one class, S-a. We therefore asked whether all four classes show both ON and OFF responses. We imaged the responses of different classes by using *Gal4* lines that target each class with either complete (S-a, S-b, I-b) or partial (I-a) specificity. Each subset of neurons showed an ON response to all four bitter compounds that were tested, and all subsets also showed prominent OFF responses to at least a subset of the bitter compounds (Figure 2A-C). Moreover, the temporal features of the response to a given compound appeared qualitatively similar across classes (Figure 2A).

**Figure 2:**
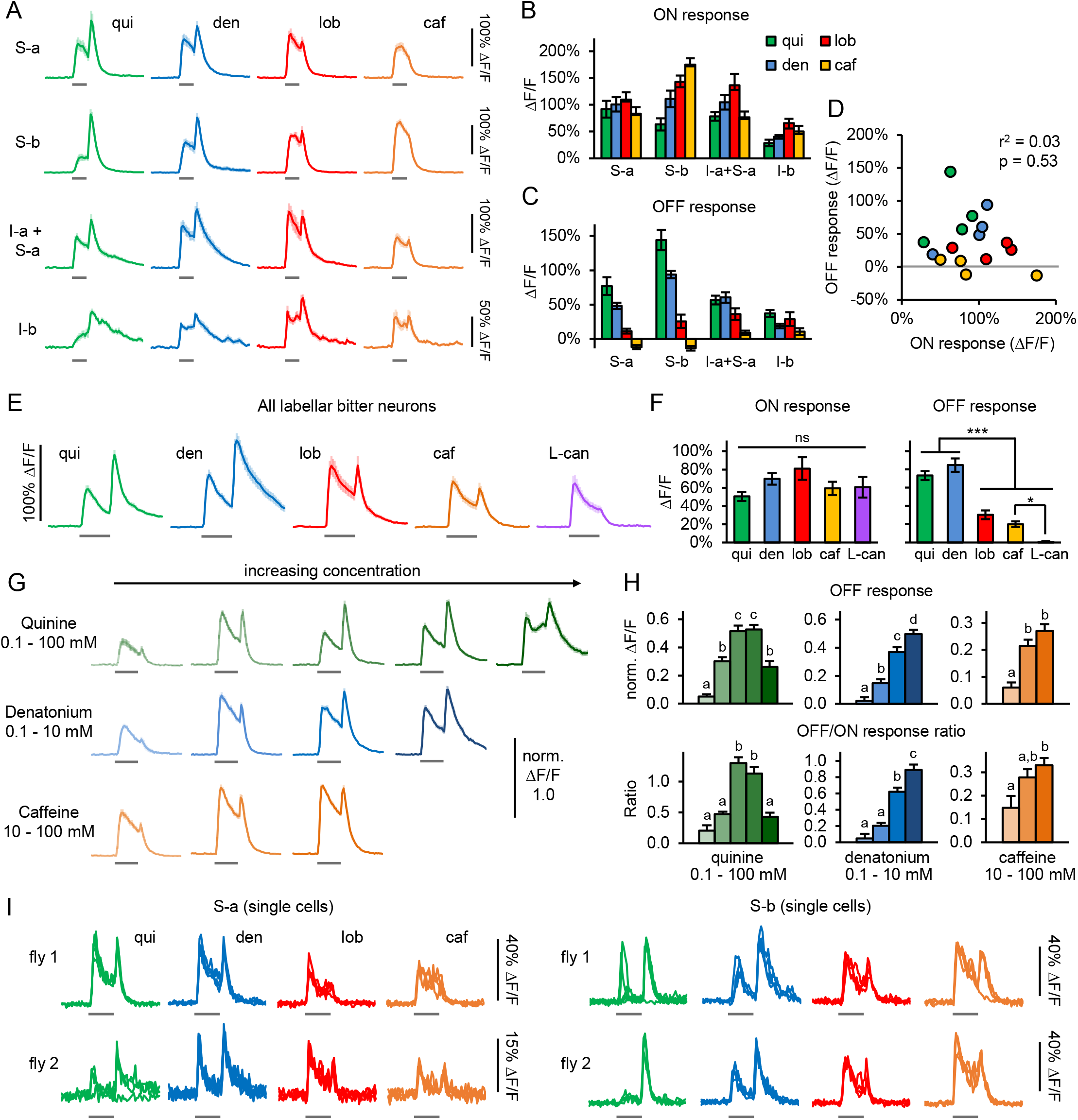
The bitter OFF response occurs in all classes of labellar bitter neurons and depends on stimulus identity and concentration. (A) Responses of different subsets of labellar bitter neurons to 4 bitter compounds (n ≥ 12 trials, 4 flies). OFF responses appear less pronounced than in other figures because the stimulus duration was 3 sec instead of 5 sec. (B-C) Peak ON (B) and OFF (C) responses for data shown in panel A. There was a significant effect of compound and class on both the ON and OFF response (p<0.0001, two-way ANOVA). (D) No correlation was observed between ON and OFF responses (r^2^ = .03, p = 0.53, Pearson’s correlation). Each point represents the average response of one neuronal subset to one compound; points are colored by compound as in panels A-C. (E) Stimulus-dependent bitter response dynamics were observed using a *Gal4* line that labels all labellar classes (n = 15 trials, 5 flies). (F) Peak ON and OFF responses for data shown in panel E. (G) Responses of S-a bitter cells show concentration-dependent dynamics (n ≥ 12 trials, 4 flies). See Methods for concentrations used. Because not all flies were tested with all concentrations, ΔF/F values were normalized to the maximum response of each fly to enable across-fly comparisons. (H) The peak OFF response (top) and OFF/ON response ratio (bottom) generally increased with concentration (data from panel G; same legend colors). (I) Individual bitter cells show both ON and OFF responses. Traces represent individual trials. To enable stochastic single-cell labeling flies had the following genotype: *hsFLP/+; Gr-Gal4, UAS-GCaMP6f/+; FRT-Gal80-FRT/UAS-NLS-RedStinger. Gal4* lines used in this figure were *Gr98d-Gal4* (S-a), *Gr22f-Gal4* (S-b), *Gr59c-Gal4* (S-a + I-a), *Gr47a-Gal4* (I-b), and *Gr33a-Gal4* (all bitter neurons). Stimulus duration was 3 sec for panels A-D and 5 sec for panels E-I. In panels F and H responses were compared by one-way ANOVA followed by Tukey’s post-tests (*p<0.05, ***p<0.001; in panel H different letters denote a significant difference at p< 0.05). See also Figure S2.

We next asked whether the OFF response differs upon exposure to different bitter stimuli. Within each subset of bitter neurons, the magnitude of the OFF response induced by different compounds varied substantially whereas each ligand elicited comparable ON responses (Figure 2A-C). At the concentrations that were tested (see Methods), quinine and denatonium induced the strongest OFF responses, lobeline induced a weaker OFF response, and caffeine induced little to no OFF response. This response profile was observed in all four classes of labellar bitter neurons. We plotted the peak ON response versus the peak OFF response for each neuronal subset in response to each compound (Figure 2D) and observed no overall correlation between the ON and OFF response (r^2^ = .03, p = 0.53, Pearson’s correlation).

We recapitulated the stimulus-dependence of the OFF response using a *Gal4* line that labels all labellar bitter neurons (*Gr33a-Gal4*). Again we observed strong OFF responses to quinine and denatonium and weaker OFF responses to lobeline and caffeine (Figure 2E-F; note that the caffeine concentration was higher than in Figure 2A-D). Interestingly, we also found that L-canavanine induced an ON response but no OFF response at all (Figure 2E-F). It is thought that different bitter compounds activate different receptors (Scott, 2018), suggesting that the response dynamics are determined by the bitter receptors themselves and are not an intrinsic property of bitter neurons. We trained a linear classifier to decode bitter identity based on these responses. Using the OFF response alone we could decode bitter identity with 61% accuracy, well above the chance level of 20% and close to the accuracy when using both the ON and OFF responses (66%), whereas using the ON response alone performed only slightly above chance (31%) (Figure S2A). In contrast, tarsal bitter responses (Figure 1E) performed poorly in decoding stimulus identity whether we used the ON response, the OFF response (which is close to zero), or both (Figure S2B). Thus bitter responses in the labellum contain much more information about bitter identity than responses in the leg, and this is largely due to the OFF response.

We next tested whether the OFF response depends on bitter concentration. The lowest bitter concentrations that were tested elicited ON responses but no OFF response (Figure 2G-H). As the bitter concentration was increased, an OFF response emerged and generally increased in magnitude (Figure 2G-H). Moreover, the ratio of the OFF to ON response also tended to increase with increasing bitter concentration (Figure 2H). Decoder analyses revealed that either the ON or OFF response alone could decode concentration above chance level but the OFF response performed much better (Figure S2C-D), indicating that the OFF response contains more information about bitter concentration.

We also examined whether the OFF response is modulated by other factors such as sex, hunger, or the presence of an appetitive tastant. Bitter neurons in male and female flies showed similar dynamics, both displaying a prominent OFF response to quinine and denatonium (Figure S2E). Bitter neurons in fed and starved flies also showed similar dynamics (Figure S2F). Finally, the addition of sucrose to bitter solutions did not modulate the dynamics of the bitter response (Figure S2G). The presence of an OFF response to sugar-bitter mixtures is important because, in nature, flies first taste a potential food source with their legs and are much more likely to sample it with their labellum if it contains an appetitive tastant such as sugar (Flood et al., 2013; Scott, 2018). Overall, these results show that bitter OFF responses are observed in all classes of labellar bitter neurons and depend on stimulus identity and concentration, but not on other factors such as sex, hunger, or the presence of sugar.

### Individual bitter sensory neurons produce both ON and OFF responses

In other systems, such as the visual system, ON and OFF responses to a particular stimulus are generated by separate neurons, affording the ability to activate different neural pathways in response to stimulus onset and offset (Hubel and Wiesel, 1961; Westheimer, 2007). In contrast, a system in which the same cells produce ON and OFF responses suggests that the same downstream pathways are activated at both times. We therefore asked whether individual bitter-sensing cells produce both ON and OFF responses.

When imaging axonal projections of labellar bitter neurons it is not possible to distinguish the responses of individual GCaMP-expressing cells. We therefore employed stochastic labeling using FLP-mediated excision of the Gal4 repressor *Gal80*, and we identified flies that contained a single labeled labellar neuron. All 15 cells that were imaged showed both ON and OFF responses to at least some bitter stimuli, and in no case did a bitter neuron exclusively show an ON or OFF response (Figure 2I). Thus individual bitter cells produce both ON and OFF responses, implying that the same downstream pathways may be activated at the onset and offset of each stimulus.

### Bitter OFF responses are generated cell-intrinsically

We tested whether the bitter OFF response is generated cell-intrinsically or through extrinsic circuit mechanisms that are likely to rely on synaptic input onto bitter sensory neurons. Although synaptic inputs onto the dendrites of labellar taste neurons have not been reported, their axon terminals may receive modulatory inputs (Chu et al., 2014; Inagaki et al., 2012; LeDue et al., 2016). Thus if the OFF response is generated by a circuit mechanism, it should not be observed in upstream compartments. We imaged bitter neuron axons in the labial nerve before they enter the brain and observed strong ON and OFF responses (Figure 3A-C), suggesting that the OFF response is generated cell-intrinsically. Nerve responses showed smaller OFF responses and less ON response adaptation than the axon terminals (Figure 3A-C), implying that some aspects of the response may be modulated by synaptic input or subcellular transformations of activity.

**Figure 3:**
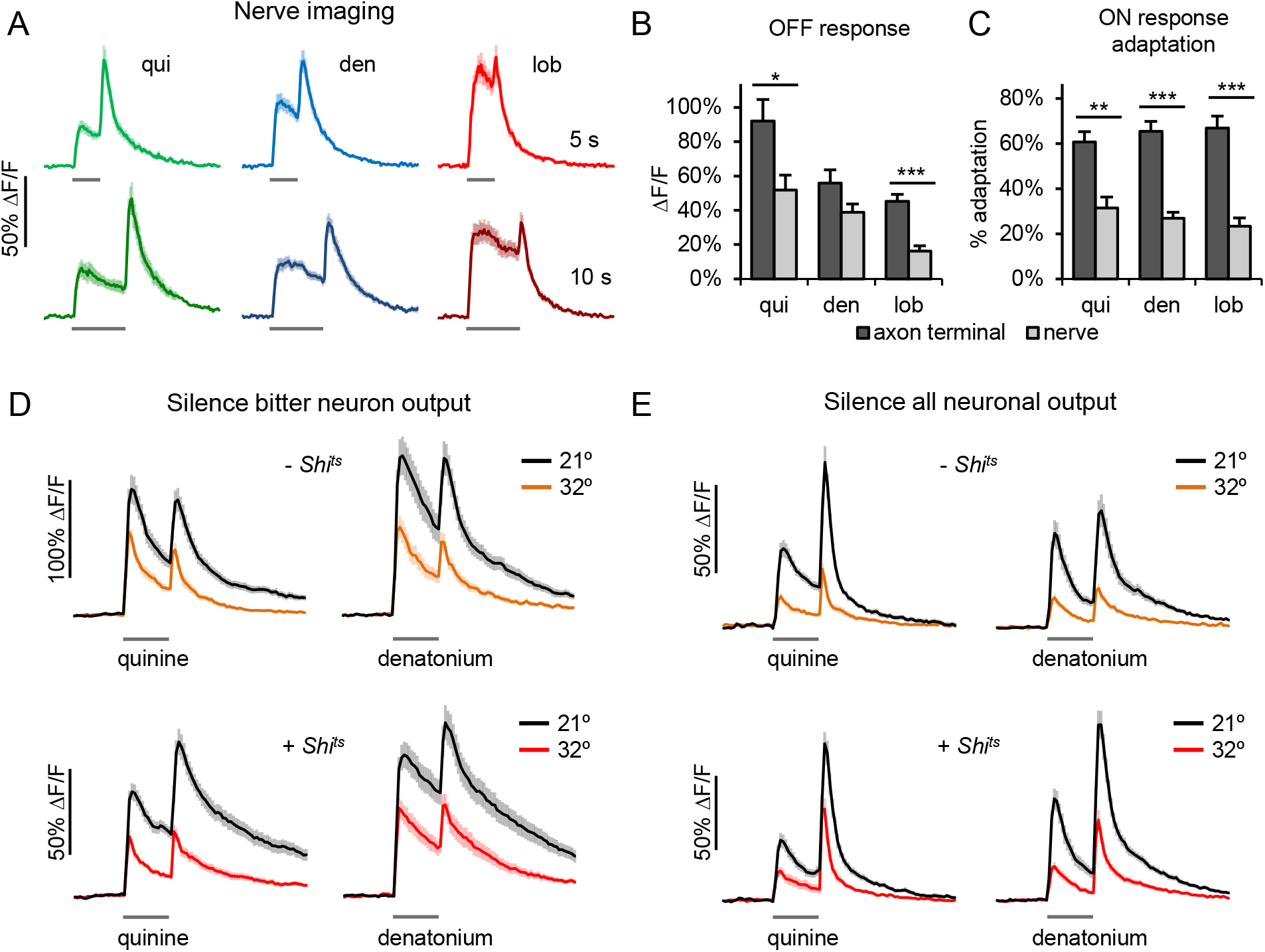
The bitter OFF response is generated cell-intrinsically. (A) Bitter OFF responses were observed in *Gr98d-Gal4*-expressing labellar bitter axons imaged in the labial nerve ∼150-200 µm upstream of the axon terminals (n ≥ 14 trials, 5 flies). Stimulation was for 5 or 10 sec. (B-C) Comparison of the peak OFF response (B) and ON response adaptation (C) between responses of labellar bitter neurons imaged in the nerve (data from panel A) or in the SEZ (data from Figure 1C) using 5 sec bitter stimulation. The nerve generally showed smaller OFF responses and less ON response adaptation than the axon terminals (*p<0.05, **p<0.01, ***p<0.001, Mann-Whitney test). (D) Silencing the synaptic output of bitter neurons did not affect their response dynamics. Control flies (“-*Shi*^*ts*^”) were *Gr33a-Gal4 > UAS-GCaMP6f* (n = 16 trials, 4 flies); experimental flies (“+ *Shi*^*ts*^”) were *Gr33a-Gal4 > UAS-GCaMP6f + UAS-Shi*^*ts*^ (n ≥ 28 trials, 7 flies). (E) Bitter OFF responses persisted when synaptic transmission was silenced pan-neuronally. Control flies (“-*Shi*^*ts*^”) were *Gr66a-lexA > lexAop-GCaMP6f, nsyb-Gal4* (n = 24 trials, 6 flies); experimental flies (“+ *Shi*^*ts*^”) were *Gr66a-lexA > lexAop-GCaMP6f, nsyb-Gal4 > UAS-Shi*^*ts*^ (n = 24 trials, 6 flies). Stimulus duration was 5 sec for panels D-E.

We next conducted experiments to silence synaptic transmission using the temperature-sensitive dynamin allele *Shibire*^*ts*^ (*Shi*^*ts*^). We prevented feedforward signaling from the bitter neurons to test the possibility of a feedback circuit mechanism that could generate the OFF response. Flies expressing *UAS-Shi*^*ts*^ in addition to *UAS-GCaMP6f* in all bitter neurons (*Gr33a-Gal4*) were compared to control flies that did not carry *UAS-Shi*^*ts*^. Both groups of flies were tested at 21°, at which *Shi*^*ts*^ is not active and synaptic transmission is unaffected, and at 32°, at which *Shi*^*ts*^ blocks synaptic transmission. We found that bitter ON and OFF responses were generally diminished at 32° in both groups of flies, apparently reflecting a suppressive effect of high temperature, but experimental and control flies did not differ in their dynamics (Figure 3D). Experimental flies continued to show prominent OFF responses at 32° despite the lack of synaptic output from bitter neurons (Figure 3D), indicating that OFF responses are not generated by a feedback circuit mechanism.

We also imaged the responses of bitter neurons (*Gr66a-lexA* driving *lexAop-GCaMP6f)* while silencing synaptic transmission in all brain neurons (*nsyb-Gal4* driving *UAS-Shi*^*ts*^). When flies of this genotype were transferred to 32° they were completely paralyzed, demonstrating the efficacy of synaptic silencing. However, bitter neurons in these flies continued to show both ON and OFF responses at 32°, and the dynamics were comparable to control flies lacking *Shi*^*ts*^ (Figure 3E). These results demonstrate that bitter OFF responses persist in the absence of synaptic transmission and are likely to result from cell-intrinsic mechanisms.

### OFF responses in bitter neurons require functional bitter receptors

We next examined whether the OFF response is generated by bitter receptors or instead results from a more general property of the sensory neuron (Figure 4A). We first tested whether an OFF response is generated following optogenetic activation of the bitter neurons using the light-activated cation channel Chrimson (Klapoetke et al., 2014). Chrimson-expressing bitter neurons were consistently activated by red light, but failed to produce an OFF response when the light was turned off (Figure 4B) Labellar bitter neurons that did not express Chrimson did not show a response to photostimulation (Figure 4B). Thus, OFF responses in bitter neurons are not a general consequence of neuronal depolarization.

**Figure 4:**
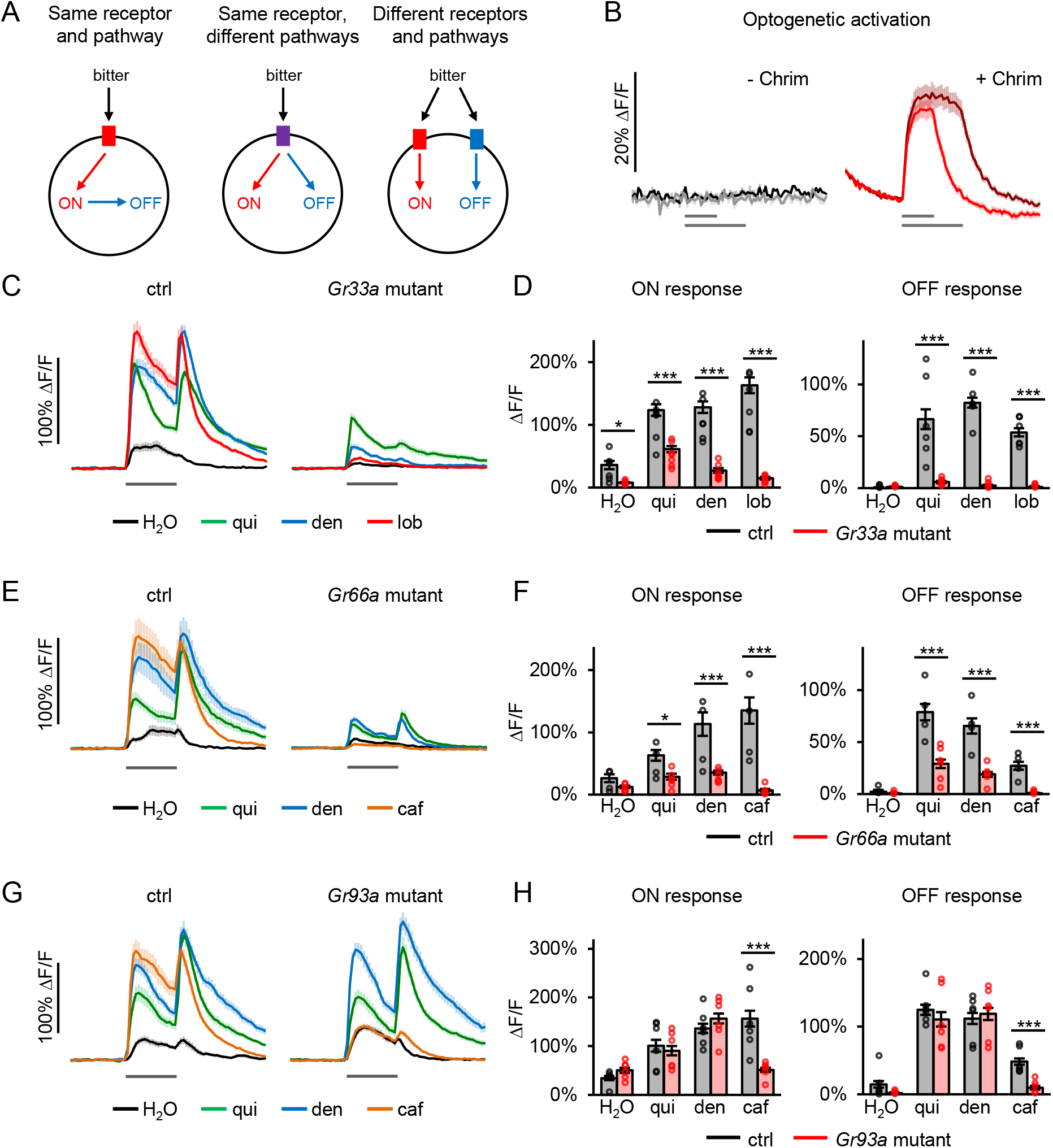
OFF responses in bitter-sensing neurons are not a general consequence of depolarization and require functional bitter receptors. (A) Models for generating the bitter OFF response. The OFF response could reflect a general cellular mechanism producing rebound excitation when the ON response is terminated (left model) or could require bitter ligands to bind specific receptors, such as canonical gustatory receptors. In the latter case, ON and OFF responses could be generated by the same receptor (center model) or two different receptors (right model). (B) Responses of *Gr98d-Gal4*-expressing labellar bitter neurons to optogenetic activation using Chrimson. Controls (“-Chrim”) were *Gr98d-Gal4 > UAS-GCaMP6f* and did not show a response to photostimulation (n = 4 trials, 2 flies). Experimentals (“+ Chrim”) were *Gr98d-Gal4 > UAS-GCaMP6f +UAS-Chrimson* and showed an ON response but no OFF response to photostimulation (n = 21 trials, 7 flies). Chrimson-expressing neurons also showed a response to the two-photon laser (925 nm) that gradually decayed during each trial. (C-D) Mutation of *Gr33a* impaired ON and OFF responses to multiple compounds. Heterozygous controls (*Gr33a-Gal4*^*mut*^ */ UAS-GCaMP6f*; n = 18 trials, 6 flies) were compared to transheterozygous mutants (*Gr33a-Gal4*^*mut*^ */ Gr33a*^*1*^, *UAS-GCaMP6f*; n = 21 trials, 7 flies). (E-F) Mutation of *Gr66a* impaired ON and OFF responses to multiple compounds. Heterozygous controls (*Gr66a-Gal4 / UAS-GCaMP6f*; *Gr66a*^*ex83*^ */ +*; n = 12 trials, 4 flies) were compared to homozygous mutants (*Gr66a-Gal4 / UAS-GCaMP6f*; *Gr66a*^*ex83*^; n = 18 trials, 6 flies). (G-H) Mutation of *Gr93a* impaired ON and OFF responses to caffeine but not to other bitter compounds. Heterozygous controls (*Gr66a-Gal4 / UAS-GCaMP6f*; *Gr93a*^*3*^ */ +*; n ≥ 17 trials, 6 flies) were compared to homozygous mutants (*Gr66a-Gal4 / UAS-GCaMP6f*; *Gr93a*^*3*^; n = 18 trials, 6 flies). Data points in panels D, F, and H represent average responses for individual flies; bars represent average responses across all flies and trials. All bitter stimuli were delivered for 5 sec. *p<0.05, ***p<0.001, two-way ANOVA followed by Bonferroni post-tests.

We next asked whether known bitter receptors are required for the OFF response. Each labellar bitter neuron expresses between 6 and 28 bitter receptor subunits that are thought to form heteromeric receptor complexes (Dweck and Carlson, 2020; Lee et al., 2009; Shim et al., 2015; Weiss et al., 2011). We first examined bitter responses in flies carrying a mutation in *Gr33a*, a putative bitter co-receptor that is broadly required for bitter responses (Dweck and Carlson, 2020; Moon et al., 2009). *Gr33a* mutant flies showed impaired ON and OFF responses to all three bitter compounds tested, with some responses affected more severely than others (Figure 4C-D). We also tested flies carrying a mutation in *Gr66a*, another broadly expressed bitter receptor that has been implicated in the response to caffeine (Dweck and Carlson, 2020; Moon et al., 2006). *Gr66a* mutants showed a complete deficit in the ON and OFF response to caffeine as well as strong impairments in responses to quinine and denatonium (Figure 4E-F). Finally, we tested flies carrying a mutation in *Gr93a*, a receptor required for detecting caffeine but not other bitter compounds (Lee et al., 2009). In *Gr93a* mutant flies the ON and OFF responses to caffeine were reduced to the level of the response to water, whereas responses to quinine and denatonium were not affected (Figure 4G-H). Thus different receptors mediate OFF responses to different compounds and the same receptor subunit mediates the ON and OFF response to a given compound, but it is not clear whether Gr93a does so by functioning in a single receptor complex (Figure 4A, center model) or two different complexes (Figure 4A, right model). Together, these results show that canonical bitter receptors are required for generating bitter OFF responses.

### Bitter ON and OFF responses are propagated to downstream dopaminergic neurons

We next examined whether ON and OFF responses can be observed in neurons downstream of bitter sensory cells. Bitter signals are transmitted to the PPL1 subset of dopaminergic neurons (DANs) that innervate the mushroom body, the primary learning and memory center of the fly (Kim et al., 2017; Kirkhart and Scott, 2015). We first confirmed that four of the five PPL1 DANs innervating the mushroom body responded to labellar bitter stimulation (Figure 5 and Figure S3), as previously reported (Kirkhart and Scott, 2015). The DANs innervating the γ2α’1, α3, and γ1 compartments of the mushroom body showed consistent ON and OFF responses to labellar stimulation with quinine and denatonium (Figure 5 and S3). The α2 α’2 DAN showed weaker and less consistent ON and OFF responses (Figure S3A) and the α’3 DAN did not respond to bitter (data not shown). In contrast, applying these compounds to the leg elicited a strong ON response in the PPL1 DANs but no OFF response (Figure 5B-C). PPL1 DANs also showed similar ligand-dependent dynamics (Figure 5D) as labellar bitter sensory neurons (Figure 2E). Bitter response dynamics in the DANs were not modulated by hunger or the presence of sugar (Figure S3C-D), similar to our results in sensory neurons (Figure S2F-G). Thus, both ON and OFF responses of bitter sensory neurons are propagated to the PPL1 DANs.

**Figure 5:**
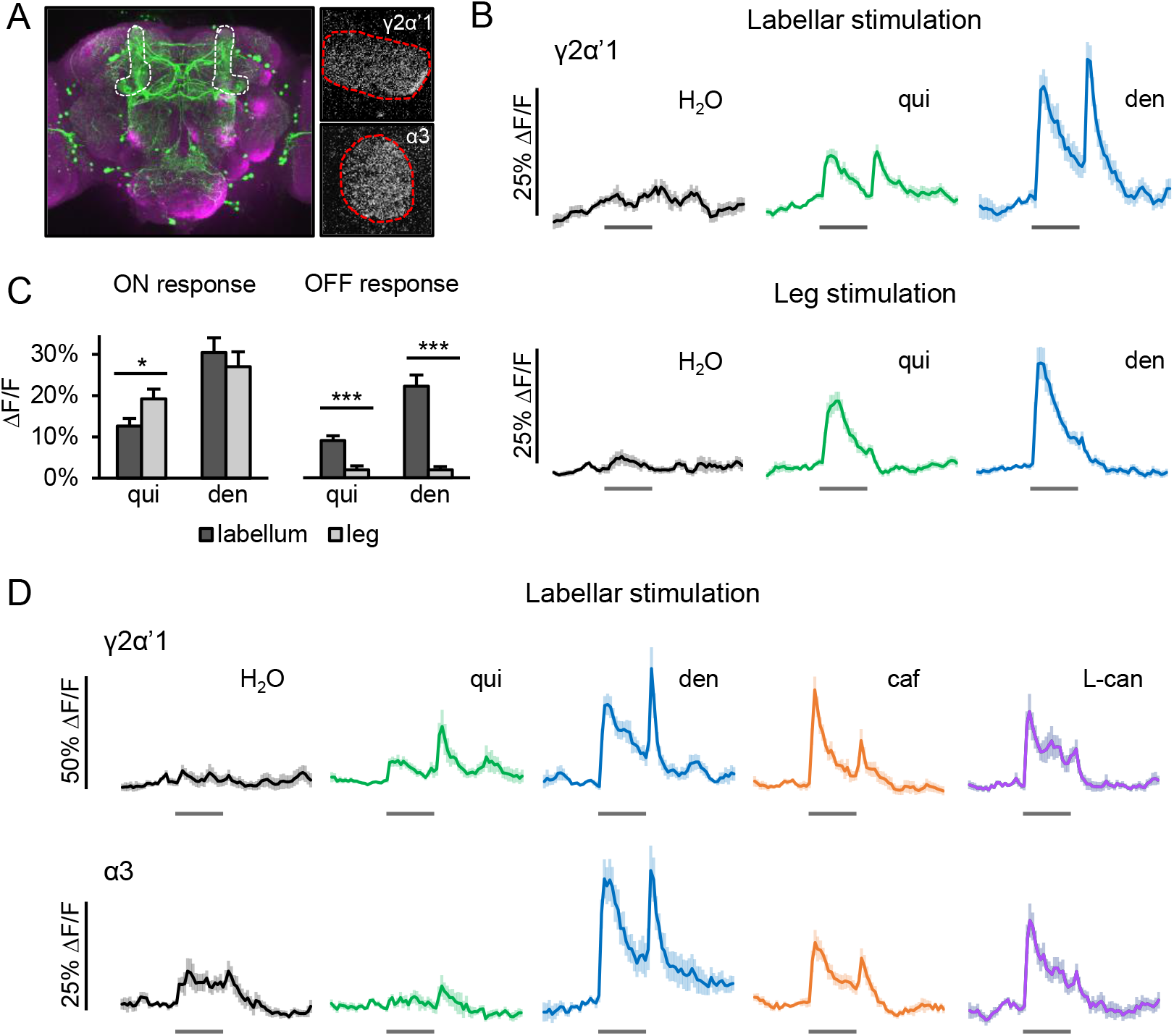
Bitter ON and OFF responses are propagated to PPL1 DANs. (A) PPL1 DANs were targeted by *TH-Gal4*. Left: fixed and stained brain showing *TH-Gal4* expression pattern. The vertical lobes of the mushroom body containing axons of PPL1 DANs are circled. Right: example frames from live GCaMP imaging of γ2α’1 or α3 DAN axons in the mushroom body lobes (maximum projections of 10-20 frames in t-series). (B) The γ2α’1 DAN showed ON and OFF responses to bitter stimulation of the labellum (top row; n ≥ 23 trials, 7 cells, 4 flies), but showed only an ON response to bitter stimulation of the leg (bottom row; n ≥ 26 trials, 3-6 flies). (C) Peak ON and OFF responses of the γ2α’1 DAN to labellar and leg stimulation (*p<0.05, ***p<0.001, Mann-Whitney test). (D) The γ2α’1 and α3 DANs showed ligand-dependent dynamics to labellar bitter stimulation (n = 12 trials, 4 flies). All bitter stimuli were delivered for 5 sec. See also Figure S3.

### Pairing odor and bitter induces unexpected effects on MBON plasticity

The timing of PPL1 DAN activity influences synaptic plasticity during aversive olfactory learning, in which an odor (the conditioned stimulus, CS) is paired with a punishment (the unconditioned stimulus, US). Odor-responsive Kenyon cells (KCs) synapse onto mushroom body output neurons (MBONs), and US-encoding PPL1 DANs modulate the strength of synapses onto MBONs that promote attraction (Figure 6A) (Aso et al., 2014a; Aso et al., 2014b; Owald and Waddell, 2015). During aversive conditioning, the simultaneous presentation of an odor with a punishment leads to coincident activation of KCs and PPL1 DANs, which induces depression at active KC-MBON synapses and results in odor avoidance (Cohn et al., 2015; Hige et al., 2015). In contrast, DAN activity prior to the odor or in the absence of odor induces MBON facilitation (Berry et al., 2018; Cohn et al., 2015; Handler et al., 2019). Thus the timing of DAN activity is critical in determining plasticity, and usually this timing coincides with the presence of a US. However, using labellar bitter stimulation as a US leads to both an ON and OFF response in the DANs (Figure 5 and S3), and the OFF response is not coincident with bitter. If bitter is paired with odor using a typical aversive learning protocol, in which odor predicts bitter onset and terminates at or before bitter offset, the odor will overlap with the bitter ON response but not the OFF response (Figure 6B, top). Conversely, if bitter is presented first and odor predicts bitter removal (Figure 6B, bottom), representing a typical “relief learning” paradigm (Tanimoto et al., 2004), then odor will overlap with the bitter OFF response but not the ON response. In each case one peak of DAN activity coincides with odor while the other does not, suggesting that they may drive plasticity in opposite directions. The bitter ON and OFF responses in the DAN thus raise an intriguing question of what type of plasticity will be observed when bitter is used as a reinforcement cue for learning.

**Figure 6:**
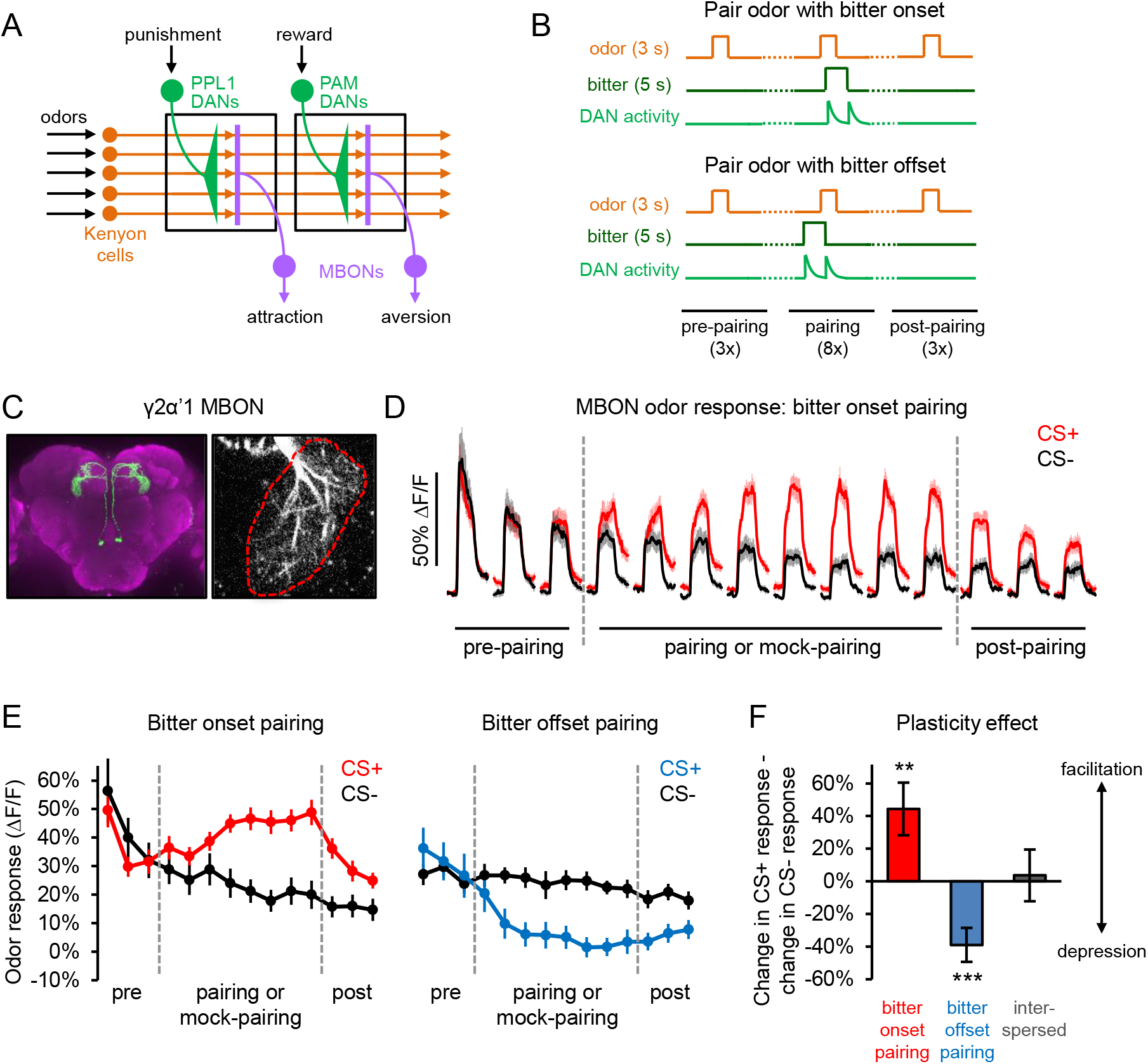
Odor-bitter pairing induces unexpected effects on MBON plasticity. (A) Schematic of the mushroom body circuit. During aversive olfactory conditioning, PPL1 DANs induce plasticity at synapses onto attraction-promoting MBONs, and the timing of their activity relative to the odor determines the direction of plasticity. (B) Schematic of odor-bitter pairing experiments. In separate experiments, the CS+ odor was paired with bitter onset (overlapping with the bitter ON response) or bitter offset (overlapping with the bitter OFF response). The odor interpulse interval (IPI) was 10 sec. Each fly was also tested with a CS-odor, which was delivered with the same timing as the CS+ but was not paired with bitter (“mock-pairing”). Each fly was tested with three interleaved CS+ and CS-blocks. (C) Images of the γ2α’1 MBON labeled with the split *Gal4* line *MB077C*. Left: fixed and stained brain. Right: example frame from live GCaMP imaging of γ2α’1 MBON dendrites. (D) GCaMP responses of the γ2α’1 MBON to CS+ and CS-odors before, during, and after pairing with bitter onset (for the CS+) or mock-pairing (for the CS-; n = 5 flies, 15 blocks). The GCaMP signal between odor pulses is not shown. (E) Average odor response of the γ2α’1 MBON during bitter onset (n = 5 flies, 15 blocks) and bitter offset (n = 7 flies, 21 blocks) pairing protocols. For the bitter offset pairing graph, we only averaged the initial odor response before the vacuum turned on because the vacuum induced a change in airflow that led to MBON activation (see Figure S4D). However, the entire odor response was averaged for analyses that do not involve consideration of paired trials, such as the analysis in panel F. (F) MBON plasticity was quantified as the change in CS+ response after pairing minus the change in CS-response. Pairing odor with bitter onset versus bitter offset induced opposite plasticity effects, whereas interspersing odor and bitter stimuli did not induce significant plasticity (n = 5 flies, 15 blocks). Error bars represent standard deviation obtained by bootstrapping, and p-values were obtained by permutation testing (**p<0.01, ***p<0.001). See also Figure S4.

We first paired odor with bitter onset, representing a typical aversive learning protocol (Figure 6B, top). Odor-bitter pairing was repeated 8 times with a 10 sec inter-odor interval. We examined whether this induced MBON plasticity by comparing the response of the γ2α’1 MBON (Figure 6C) to the conditioned odor (CS+) before and after pairing. As a control, we repeated the same protocol for a different odor (the unconditioned odor, CS-) without bitter pairing (“mock-pairing”). The MBON response to the CS-decreased with repeated presentations, representing odor habituation (Figure 6D). The effect of odor-bitter pairing was quantified by calculating the percent change in the CS+ and CS-responses after pairing or mock-pairing and then taking the difference. Whereas pairing an odor with an aversive US normally induces MBON depression (Berry et al., 2018; Handler et al., 2019; Perisse et al., 2016), pairing odor with bitter instead enhanced the MBON response (Figure 6D-F; compare CS+ to CS-). This facilitation was most pronounced during odor-bitter pairing but also persisted when the odor was presented alone. We also observed that the CS+ response was enhanced on the first bitter pairing trial before any learning has occurred (Figure 6D-E), suggesting that bitter acutely enhances MBON odor responses even though bitter alone does not reliably activate the MBON (Figure S4A-B). Pairing odor with water instead of bitter did not induce MBON plasticity (Figure S4C), consistent with our observation that water does not activate the γ2α’1 DAN (Figure 5).

We next paired odor with bitter offset, representing a typical relief learning or “backward pairing” protocol in which the US precedes the CS (Figure 6B, bottom). Performing this type of conditioning with electric shock induces MBON facilitation (Handler et al., 2019), but using bitter we instead observed MBON depression (Figure 6E-F). Because bitter stimuli are removed by a vacuum line, bitter offset coincides with a change in airflow that often activated the MBON (Figure S4D). However, the vacuum stimulus was also presented in CS-blocks, and we conducted additional analyses showing that the vacuum is unlikely to drive MBON depression (Figure S4D). These results demonstrate that pairing odor with bitter onset versus bitter offset leads to opposing effects on MBON plasticity, and both effects contrast with the effects induced by other USs. No MBON plasticity was observed when odor and bitter stimuli were interspersed (Figure 6F).

### The bitter OFF response drives plasticity during repeated odor-bitter pairing

The MBON facilitation induced by pairing odor with bitter onset and the MBON depression induced by pairing odor with bitter offset both differ from previous results using other aversive USs (Berry et al., 2018; Handler et al., 2019; Perisse et al., 2016). We hypothesized that the DAN’s OFF response to bitter, which has not been reported for other USs, might account for these unexpected effects. This explanation was bolstered by a surprising observation: when we imaged the DAN’s response to repeated bitter stimulation at 10 sec intervals, similar to the timing of bitter presentation during pairing experiments, we found that the ON response strongly habituated whereas the OFF response remained strong (Figure 7A-C). The ON response decayed by more than 85% within 3 trials, whereas the OFF response decayed by less than 10% within 3 trials and by less than 20% by the 8^th^ trial. This preferential habituation of the ON response was also observed in labellar bitter sensory neurons, indicating that it arises peripherally (Figure 7D-F and Figure S5A-B).

**Figure 7:**
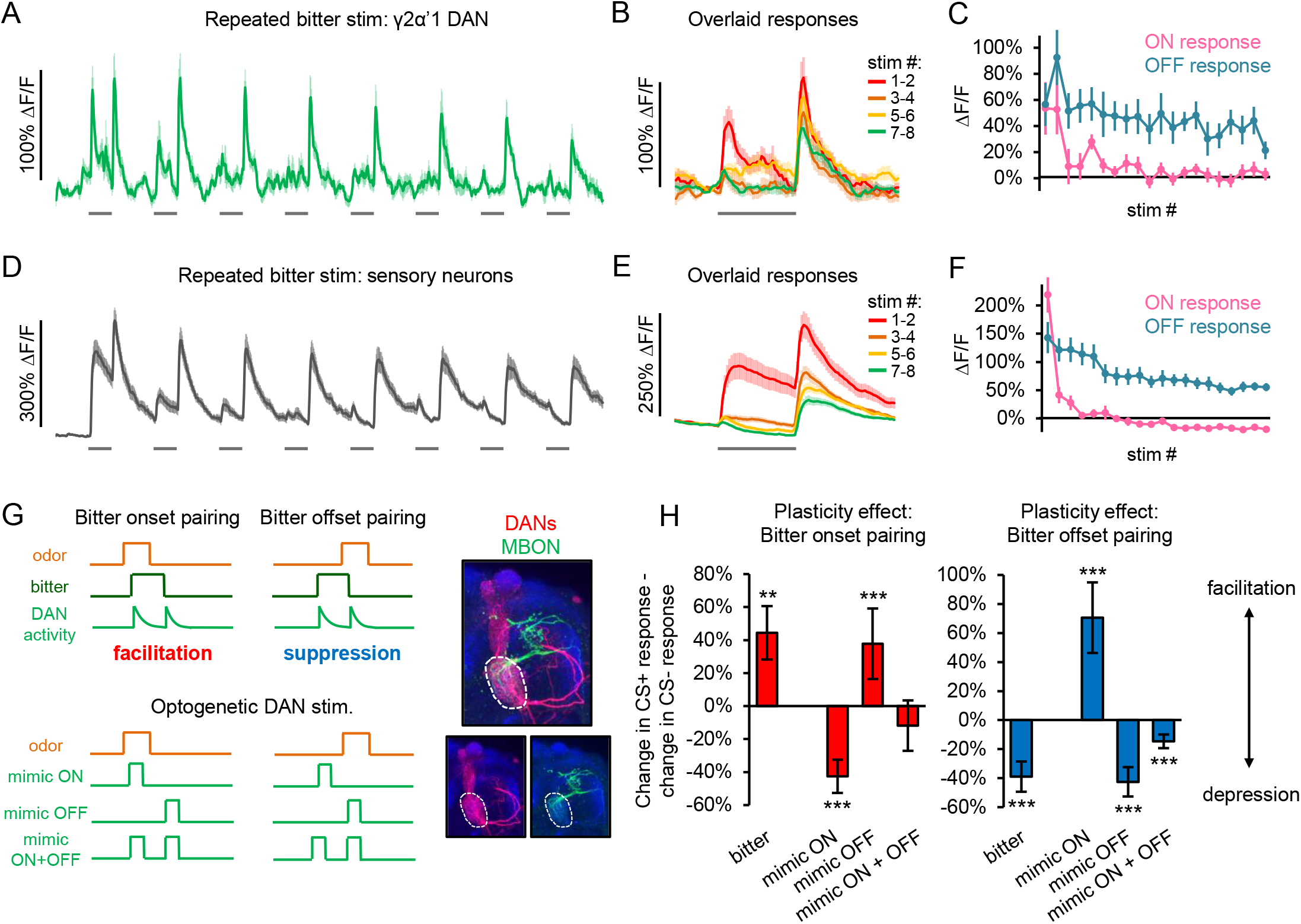
The bitter ON response rapidly habituates while the OFF response drives plasticity. (A-F) Responses of the γ2α’1 DAN (panels A-C; labeled with *TH-Gal4*, n = 9 flies) or bitter sensory neurons (panels D-F; labeled with *Gr33a-Gal4*, n = 7 flies) to repeated 5 sec denatonium stimuli delivered with a 10 sec IPI. Both sensory neurons and DANs show strong habituation of the ON response but not the OFF response. (A, D) Average GCaMP traces during the first 8 stimulations. (B, E) Average responses to different stimuli overlaid. Baseline fluorescence was calculated separately for each stimulus. (C, F) Average ON and OFF responses to each stimulus. (G-H) Odor was paired with optogenetic DAN stimulation and the effect on MBON plasticity was examined (*MB099C-splitGal4 > UAS-Chrimson, 25D01-lexA > lexAopGCaMP6f*; n = 6-8 flies, 18-24 blocks). (G) Left: DAN activation was timed to mimic the timing of the bitter ON and/or OFF response in either the bitter onset pairing or offset pairing protocols shown in Figure 6A. Right: immunostaining of fly genotype used for imaging, showing the γ2α’1 DAN (as well as the α2 α’2 DAN) labeled with *MB099C-splitGal4* and the γ2α’1 MBON labeled with *25D01-lexA*. The γ2α’1 compartment is outlined. (H) Mimicking the ON and OFF responses induced opposing effects on MBON plasticity, and only mimicking the OFF response reproduced the effects induced by bitter. Error bars represent standard deviation obtained by bootstrapping, and p-values were obtained by permutation testing. Plasticity values for “bitter” represent the same data shown in Figure 6F. See also Figure S5.

Because our pairing experiments involved pairing odor and bitter 8 times, the preferential habituation of the bitter ON response in the DAN suggests that reinforcement may be largely due to the OFF response. We tested this hypothesis by performing pairing experiments in which we replaced bitter with optogenetic stimulation of the γ2α’1 DAN. We confirmed that the DAN showed a strong response to repeated optogenetic stimulation that did not habituate over 8 trials (Figure S5C). On each pairing trial we stimulated the DAN in either a one- or two-peaked manner to mimic the timing of the bitter ON response, the OFF response, or both, and we asked which of the two peaks reproduces the effects of bitter pairing (Figure 7G-H).

We first employed photostimulation to mimic the protocol in which odor is paired with bitter onset, which induces MBON facilitation (Figure 6D-F). When we mimicked only the ON response, such that the DAN is only activated during odor presentation, we observed MBON depression (Figure 7G-H). In contrast, mimicking the timing of the OFF response, such that the DAN is activated after odor removal, resulted in MBON facilitation (Figure 7G-H). Mimicking both the ON and OFF response did not induce significant plasticity (Figure 7G-H). When these experiments were repeated in control flies that did not express Chrimson, either no plasticity or much weaker plasticity was observed (Figure S5D); weak plasticity effects may be due to red light representing an aversive US. These experiments demonstrate that when odor is paired with bitter onset, the bitter ON and OFF responses induce opposing forms of MBON plasticity. The facilitation driven by the OFF response is the dominant effect that is observed, likely due to the habituation of the ON response.

We next mimicked the protocol in which odor is paired with bitter offset, which induces MBON depression (Figure 6E-F). Optogenetically mimicking the timing of the ON response, such that the DAN is activated before odor onset, resulted in MBON facilitation (Figure 7G-H). In contrast, mimicking the timing of the OFF response, such that the DAN is activated during odor presentation, led to MBON depression (Figure 7G-H). Mimicking both the ON and OFF response led to weaker MBON depression (Figure 7G-H). Control flies that did not express Chrimson showed either no effect or much weaker plasticity for all three protocols (Figure S5D). Thus when odor is paired with bitter offset, the bitter ON and OFF responses induce opposing types of plasticity, similar to our results with bitter onset pairing, and the depression driven by the OFF response dominates. Overall, our results demonstrate that repeatedly pairing odor with either bitter onset or offset leads to the opposite effects on MBON plasticity than with other aversive USs, and these unexpected effects can be explained by the bitter OFF response.

## DISCUSSION

We found that bitter neurons in the labellum display both ON and OFF responses. In contrast, neurons activated by other tastants exhibit only an ON response. The ON and OFF responses result from cell-intrinsic mechanisms and require canonical bitter receptors. The bitter ON and OFF responses in sensory neurons are propagated downstream to PPL1 DANs and have a complex influence on synaptic plasticity during aversive learning. ON-OFF bitter cells in the *Drosophila* taste system therefore influence plasticity in neural circuits governing learning and memory.

### Bitter OFF responses in the labellum

The strong OFF response we observe in labellar bitter-sensing neurons has not been previously described in other studies of *Drosophila* bitter neurons. This may reflect differences in recording techniques or tastant delivery. Electrophysiological recordings of taste sensory neurons typically use a method that precludes neural recordings after the stimulus is removed (Dweck and Carlson, 2020; Hiroi et al., 2004; Moon et al., 2009; Weiss et al., 2011). Calcium imaging studies did not monitor the response after bitter offset (Marella et al., 2006), delivered bitter tastants through a continuous liquid flow without precise control of bitter offset (Inagaki et al., 2014), or used bitter compounds or concentrations that may not lead to a strong OFF response (Chu et al., 2014; Jaeger et al., 2018; LeDue et al., 2016). A recent study observed OFF responses in DANs stimulated with quinine but did not analyze this further (Siju et al., 2020).

The OFF response in labellar bitter neurons is ligand-specific: an OFF response is not evoked by all bitter ligands capable of eliciting an ON response (Figure 2E-F). These data suggest that the OFF response is mediated by specific bitter receptors rather than extrinsic circuit mechanisms. Moreover, an OFF response was not elicited following the generation of an ON response by optogenetic activation of bitter neurons. These findings are in accord with our observation that the OFF response persists after inhibiting synaptic transmission and argue strongly that the OFF response is mediated by a cell-intrinsic mechanism.

Bitter receptors are required for both the ON and OFF responses, and different receptors mediate the responses to different compounds (Figure 4). The ON and OFF responses to a given compound may be mediated by a single bitter receptor or distinct receptors. Bitter receptor subunits are thought to form heteromeric complexes with different ligand specificities (Dweck and Carlson, 2020; Shim et al., 2015). Deletion of the broadly required subunits *Gr33a* or *Gr66a* as well as the more specific subunit *Gr93a*, required for caffeine detection, results in the loss of both the ON and OFF response. This suggests that a single receptor may generate both an ON and OFF response. In one model, ligand binding activates the receptor, which then adopts a closed, inactivated state (Ulbricht, 2005). Release of the ligand may briefly relieve inactivation, opening the channel to produce an OFF response. Channel inactivation would also explain the strong adaptation observed for the ON response in labellar bitter neurons (Figure 1). Alternatively, release of the ligand may initiate a signaling cascade that opens a different cation channel to generate the OFF response.

### Bitter ON and OFF responses influence synaptic plasticity

The bitter ON and OFF responses in the sensory neurons are propagated to PPL1 DANs. Pairing an odor CS with bitter onset or offset as a US had a complex influence on MBON plasticity. Pairing of odor with bitter onset will result in a peak of DAN activity coincident with odor and a second peak following odor; pairing of odor with bitter offset will result in a peak of DAN activity prior to odor and a second peak coincident with odor (Figure 6B). Exposure of MBONs to multiple timed DAN activation events per odor presentation is experimentally unprecedented, and predictions are not intuitive. We found that repeated pairing of odor with bitter onset led to MBON facilitation, whereas pairing odor with bitter offset led to MBON depression. Both of these effects contrast with the sign of MBON plasticity induced by other aversive USs, such as electric shock (Berry et al., 2018; Handler et al., 2019; Perisse et al., 2016). Perhaps the simplest explanation for the plasticity effects we observe is that the bitter ON response rapidly habituates, leaving the OFF response as the dominant reinforcement signal. As a consequence, pairing odor with bitter offset results in a dominant peak coincident with odor, and the MBON suppression we observe is consistent with known timing rules (Berry et al., 2018; Cohn et al., 2015; Handler et al., 2019; Hige et al., 2015). In contrast, pairing of odor with bitter onset results in a dominant peak following the odor. The MBON facilitation we observe contrasts with a recent study showing that DAN stimulation just after an odor presentation causes MBON suppression, though DAN stimulation before the odor causes facilitation (Handler et al., 2019). This difference may reflect the fact that we performed repeated pairings with a short inter-odor interval of only 10 sec, so we cannot distinguish whether the plasticity we observed is due to DAN activity preceding or following the odor.

Because the MBON plasticity effects induced by odor-bitter pairing have the opposite sign as those induced by other USs, the learned behavior may also be reversed. This would imply that flies show odor attraction after pairing with bitter onset and odor aversion after pairing with bitter offset. Both of these effects seem ethologically counterintuitive, and it is possible that other pathways override MBON plasticity to promote adaptive behavior.

### Ethological relevance of bitter ON and OFF responses

One key question is why bitter neurons in different organs show different dynamics. Different taste organs experience bitter in different contexts and likely have different behavioral roles. Taste neurons in the legs are activated when a fly stands or walks on a substrate, and they mediate locomotor attraction to sugar and aversion to bitter (Joseph and Heberlein, 2012; Thoma et al., 2016). Tarsal bitter responses, which show a sustained ON response and no OFF response, may therefore be adapted for continuously reading out the bitterness of the substrate to mediate locomotor repulsion.

Labellar taste neurons are activated when the fly chooses to extend its proboscis to sample the substrate or ingest food (Flood et al., 2013; Yang et al., 2008). Feeding bouts are typically short, with individual labellar contacts lasting ∼0.1 to 3-4 sec depending on food texture and content (Itskov et al., 2014; Ro et al., 2014). The OFF response would occur immediately after feeding or sampling.

The bitter OFF response may be behaviorally important during longer bouts of feeding. Bitter elicits robust aversion and continued feeding in the presence of bitter is unlikely to occur unless the substrate also contains an appetitive tastant such as sucrose. A bitter-sugar mixture may create a competitive scenario during which the aversive drive, represented by the bitter ON response, undergoes significant adaptation. Within a few seconds the fly may not know it is feeding in the presence of an aversive cue. Adaptation of the bitter ON response may enable appetitive pathways to more readily override bitter-induced aversion, permitting the fly to continue feeding. A similar logic may explain why the bitter ON response habituates during repeated contacts with bitter. The fly may be choosing to feed despite the presence of bitter, and the bitter ON response may habituate to allow for this override. The OFF response may inform the fly of the presence of bitter once feeding has terminated, and this signal could be used to suppress further feeding bouts or promote locomotion to a better food source. The OFF response might also serve as a second US, in addition to the ON response, that reinforces aversive learning in an environment rich in bitter. In this manner, the bitter OFF response may elicit adaptive behaviors or reinforce memory following the consumption of aversive bitter stimuli.

## ACKNOWLEDGEMENTS

We thank Daisuke Hattori for MATLAB code and assistance with calcium imaging; Chris Rodgers and Barbara Noro for advice and comments; members of the Axel laboratory for helpful feedback and suggestions; Adriana Nemes, Phyllis Kisloff, Miriam Gutierrez, and Clayton Eccard for general laboratory and administrative support; and John Carlson, Ulrike Heberlein, Craig Montell, Kristin Scott, Janelia Research Campus, and the Bloomington Drosophila Stock Center (BDSC) for generously providing fly strains.

## AUTHOR CONTRIBUTIONS

A.V.D conceived the project and performed all calcium imaging experiments and data analysis. J.U.D. performed immunohistochemistry and confocal imaging and provided technical assistance in other areas. B.S. assisted with some MBON imaging experiments. A.V.D. and R.A. wrote the manuscript.

## MATERIALS AND METHODS

### Fly strains and maintenance

Flies were reared at 25°C and 70% relative humidity on standard cornmeal food. Experiments were performed on 1-2 week-old flies. All flies tested were females unless otherwise specified. Flies used for optogenetic experiments were maintained in constant darkness and fed on food containing 1 mM all trans-retinal for 3-5 days prior to testing. For starvation experiments, flies were food-deprived with water (using a wet piece of Kimwipe) for the specified amount of time before testing. For FLP-out experiments to image single cells, we used a heat shock-inducible FLP recombinase that induces low levels of recombination in the absence of heat shock.

Genotypes for each experiment are described below, and fly strains and their sources are described in the supplemental Key Resources Table.

### Fly genotypes used in each figure

Figure 1:

- *Gr98d-Gal4/UAS-GCaMP6f*^*p40*^ (bitter-sensing neurons)
- *Gr64f-Gal4/UAS-GCaMP6f*^*p40*^ (sugar-sensing neurons)
- *ppk28-Gal4*^*7*.*1*^*/UAS-GCaMP6f*^*p40*^ (osmolarity-sensing neurons)

Figure 2:

- *Gr98d-Gal4/UAS-GCaMP6f*^*p40*^ (S-a bitter neurons)
- *Gr22f-Gal4/UAS-GCaMP6f*^*p40*^ (S-b bitter neurons)
- *UAS-GCaMP6f*^*p40*^*/+; Gr59c-Gal4/+* (S-a and I-a bitter neurons)
- *Gr47a-Gal4/UAS-GCaMP6f*^*p40*^ (I-b bitter neurons)
- *hsFLP/+; Gr98d-Gal4, UAS-GCaMP6f*^*p40*^*/+; FRT-Gal80-FRT/UAS-NLS-RedStinger* (S-a single cells)
- *hsFLP/+; Gr22f-Gal4, UAS-GCaMP6f*^*p40*^*/+; FRT-Gal80-FRT/UAS-NLS-RedStinger* (S-b single cells)

Figure 3:

- *Gr98d-Gal4/UAS-GCaMP6f*^*p40*^ (nerve imaging)
- *Gr33a-Gal4, UAS-GCaMP6f*^*p40*^*/+; UAS-Shi*^*ts*^*/ +* (silencing bitter output, experimental flies)
- *Gr33a-Gal4, UAS-GCaMP6f*^*p40*^*/+* (silencing bitter output, control flies)
- *Gr66a-lexA/lexAop-GCaMP6f; nsyb-Gal4*^*2-1*^*/UAS-Shi*^*ts*^ (silencing pan-neuronal output, experimental flies)
- *Gr66a-lexA/lexAop-GCaMP6f; nsyb-Gal4*^*2-1*^*/+* (silencing pan-neuronal output, control flies)

Figure 4:

- *Gr98d-Gal4/UAS-GCaMP6f*^*p40*^; *UAS-Chrimson-TdT*^*VK5*^*/+* (optogenetic activation, experimental flies)
- *Gr98d-Gal4/UAS-GCaMP6f*^*p40*^ (optogenetic activation, control flies)
- *Gr33a-Gal4/Gr33a*^*1*^, *UAS-GCaMP6f*^*p40*^ (*Gr33a* mutant; note that the *Gal4* insertion is also a *Gr33a* mutation)
- *Gr33a-Gal4/UAS-GCaMP6f*^*p40*^ (*Gr33a* heterozygous control; note that the *Gal4* insertion is also a *Gr33a* mutation)
- *Gr66a-Gal4/UAS-GCaMP6f*^*p40*^; *Gr66a*^*ex83*^ (*Gr66a* mutant)
- *Gr66a-Gal4/UAS-GCaMP6f*^*p40*^; *Gr66a*^*ex83*^*/+* (*Gr66a* heterozygous control)
- *Gr66a-Gal4/UAS-GCaMP6f*^*p40*^; *Gr93a*^*3*^ (*Gr93a* mutant)
- *Gr66a-Gal4/UAS-GCaMP6f*^*p40*^; *Gr93a*^*3*^*/+* (*Gr93a* heterozygous control)

Figure 5:

- *UAS-GCaMP6f*^*p40*^*/+; TH-Gal4/+* (PPL1 DANs)

Figure 6 and Figure S4:

- *UAS-GCaMP6f*^*p40*^*/+; MB077C-splitGal4/+* (γ2α’1 MBON)

Figure 7:

- *UAS-GCaMP6f*^*p40*^*/+; TH-Gal4/+* (γ2α’1 DAN)
- *Gr33a-Gal4/UAS-GCaMP6f*^*p40*^ (bitter sensory neurons)
- *25D01-lexA/lexAop-GCaMP6F*^*p5*^; *MB099C/UAS-Chrimson-TdT*^*VK5*^ (optogenetic plasticity experiments, experimental flies)
- *25D01-lexA/lexAop-GCaMP6F*^*p5*^; *MB099C/UAS-TdT*^*VK5*^ (line for co-staining *MB099C* and*25D01-lexA* expression)

Figure S1:

- *Gr33a-Gal4/UAS-GCaMP6f*^*p40*^

Figure S2:

- *Gr33a-Gal4/UAS-GCaMP6f*^*p40*^ (male vs. female flies)
- *Gr66a-Gal4/UAS-GCaMP6f*^*p40*^ (fed vs. starved flies)
- *Gr98d-Gal4/UAS-GCaMP6f*^*p40*^ (testing bitter-sugar mixtures)

Figure S3:

- *UAS-GCaMP6f*^*p40*^*/+; TH-Gal4/+* (α2 α’2 and γ2α’1 DANs)
- *UAS-GCaMP6f*^*p40*^*/+; MB320C-splitGal4/+* (γ1 DAN)

Figure S5:

- *UAS-GCaMP6f*^*p40*^*/+; TH-Gal4/+* (γ2α’1 DAN)
- *Gr33a-Gal4/UAS-GCaMP6f*^*p40*^ (bitter sensory neurons)
- *Gr64f-Gal4/UAS-GCaMP6f*^*p40*^ (sugar sensory neurons)
- *UAS-GCaMP6F/+ ; MB099C/UAS-Chrimson-TdT*^*VK5*^ (validation of DAN optogenetic activation, “+ Chrim”)
- *UAS-GCaMP6F/+ ; MB099C/UAS-TdT*^*VK5*^ (validation of DAN optogenetic activation, “-Chrim”)
- *25D01-lexA/lexAop-GCaMP6f*^*p5*^; *MB099C/+* (optogenetic plasticity experiments, control flies)

### Taste stimulation and calcium imaging

Tastants were delivered to the labellum or foreleg as previously described (Devineni et al., 2019) via a custom-built solenoid pinch valve system controlled by MATLAB software via a data acquisition device (Measurement Computing). Pinch valves were opened briefly (∼10 ms) to create a small liquid drop at the end of a 5 µL glass capillary, positioned such that the drop would make contact with the labellum or leg. Tastants were removed after a fixed duration by a vacuum line controlled by a solenoid pinch valve. Proper taste delivery was monitored using a side-mounted camera (Veho VMS-004), which allowed for visualization of the fly and tastant capillary using the light from the imaging laser. At least three trials of each stimulus were given. Other than experiments explicitly using repeated taste stimulation, at least one minute rest was given between trials to avoid habituation.

Unless otherwise specified, the following concentrations of tastants were used: 10 mM quinine; 10 mM denatonium; 1 mM lobeline; 10 mM (Figure 2A-D), 50 mM (Figure 4E-H), or 100 mM (Figure 2E-F, 2I, 5D, and S5B) caffeine; 25 mM L-canavanine; and 100 mM (Figure S2, S3, and S5B) or 500 mM (Figure 1) sucrose. To test the effect of varying concentration (Figure 2G-H) we used the following concentrations: quinine at 0.1 mM, 1 mM, 5 mM, 10 mM, and 100 mM; denatonium at 0.1 mM, 1 mM, 5 mM, and 10 mM; and caffeine at 10 mM, 50 mM, and 100 mM. L-canavanine was also tested at 50 mM and 100 mM and no OFF response was observed (data not shown), similar to the response at 25 mM.

Flies were imaged using previously described protocols (Devineni et al., 2019). Flies were first taped on their backs to a piece of clear tape in an imaging chamber. For proboscis taste stimulation, fine strands of tape were used to restrain the legs, secure the head, and immobilize the proboscis in an extended position. For leg taste stimulation, the two forelegs were immobilized using tape and parafilm with the distal segment exposed. To image sensory neuron projections in the SEZ, an imaging window was cut on the anterior surface of the head, the antennae were removed, and the esophagus was cut in order to visualize the SEZ clearly. The labial nerve was imaged ∼150-200 µm upstream of the axon terminals through a larger window. Initial experiments imaging the PPL1 DANs used a similar preparation as for SEZ imaging, but we later modified this preparation by moving the imaging window more dorsally to exclude the antennae and leave them intact. All MBON imaging experiments used the new preparation. We generally imaged the dendritic arbors of the γ2α’1 MBON within the mushroom body lobe. However, for optogenetic experiments in which the DAN was activated, we imaged the dendritic shaft of the γ2α’1 MBON just adjacent to the lobe because otherwise the DAN’s expression of *UAS-Chrimson-TdT* bled through into the GCaMP imaging channel. Imaging experiments were performed in modified artificial hemolymph in which 15 mM ribose is substituted for sucrose and trehalose (Marella et al., 2006; Wang et al., 2003).

Calcium imaging experiments were performed using a two-photon laser scanning microscope (Ultima, Bruker) equipped with an ultra-fast Ti:S laser (Chameleon Vision, Coherent) that is modulated by pockel cells (Conoptics). Emitted photons were collected with a GaAsP photodiode detector (Hamamatsu) through a 60X water-immersion objective (Olympus). Images were acquired using the microscope software (PrairieView, Bruker). A single plane through the widest and/or brightest area of axonal or dendritic projections was chosen for imaging. Images were acquired at 925 nm at a resolution of 256 by 256 pixels. The scanning rate was 3-4 Hz for most experiments in which sensory neurons or DANs were imaged without repeated taste stimulation (Figures 1-5). The scanning rate was increased to 6-7 Hz for MBON imaging and experiments using repeated taste stimulation (Figures 6-7).

### Single cell labeling

In order to label individual bitter neurons, we used FLP-mediated excision of *Gal80* to sparsen bitter *Gal4* expression patterns. In a small, random subset of *Gal4*-expressing neurons, *Gal80* is excised and allows Gal4 to drive expression of *UAS-GCaMP* as well as the nuclear red marker *UAS-NLS-Redstinger*. We screened hundreds of flies to identify those that contained exactly one labeled labellar neuron based on expression of Redstinger, and we then imaged its axonal projections.

### Shibire silencing

Imaging experiments involving Shi^ts^ were performed similarly to previously described protocols (Hattori et al., 2017). The imaging chamber was held in a temperature-controlled platform (Warner Instruments) mounted on the microscope stage. Flies were prepared and initially imaged at room temperature (∼21°). The saline was then exchanged with pre-heated saline at 32°. The temperature was constantly monitored using a small thermistor placed in the saline immediately next to the microscope objective and was maintained close to 32° by the heating system. After imaging at 32°, flies were returned to room temperature by turning off the heating system and exchanging the warm saline for room temperature saline. Post-heating responses at room temperature were similar to those recorded before heating (data not shown)

### Optogenetic stimulation

Optogenetic stimulation during calcium imaging was delivered as previously described (Hattori et al., 2017). Red light was delivered using a 617 nm LED (Luxeon Star) with an attached lens. Light onset and offset were triggered using the imaging software’s voltage output function through an LED controller (BuckPuck 700 mA, Luxeon Star). A high-speed shutter was closed during light delivery to protect the GaAsP detector, so images were only acquired on frames when the light was off and light delivery was timed to coincide with specific frames. For optogenetic activation and imaging of bitter sensory neurons (Figure 4B), light was turned on for every third frame: 269 ms light on, 597 ms light off, repeated 5 or 10 times for a total stimulus duration of ∼4 sec or ∼8 sec, respectively. Calcium responses were linearly interpolated during these light stimulation periods. For optogenetic activation of DANs (Figure 7H and S5C), each light stimulus comprised a 50 ms light pulse repeated 16 times with an interpulse interval of 100 ms, for a total duration of 2.3 sec.

### Odor-bitter pairing

A 3 sec odor stimulus (the conditioned odor, CS+) was turned on 0.5 sec prior to either the onset (“bitter onset pairing”) or offset (“bitter offset pairing”) of a 5 sec bitter stimulus. In each CS+ block the odor was presented 3 times alone (“pre-pairing”), then paired 8 times with bitter (“pairing”), then again presented 3 times alone (“post-pairing”). The inter-odor interval was 10 sec. CS-blocks were presented in the same way but without bitter delivery. The vacuum that removes the bitter stimulus was still presented at the same time during CS-blocks. The CS+ and CS-blocks were repeated 3 times in each fly with an interval of several minutes between blocks. The odors used were 3-octanol (OCT) and 4-methylcyclohexanol (MCH). Which odor was used as the CS+ or CS-was counterbalanced across flies. The order of CS+ and CS-blocks was also counterbalanced, i.e. half the flies experienced the CS+ block first and the other experienced the CS-block first.

Odors were delivered as previously described (Hattori et al., 2017) using a computer-controlled olfactometer (Island Motion). Odors were diluted in mineral oil at 1:250 for MCH and 1:333 for OCT. 500 µL of each odor solution was pipetted onto a Whatman syringe filter and inserted into an odor mixing manifold. An odorized air stream was created by flowing air at 500 mL/min through the manifold, with valves controlling which odor filter received the air stream. This odorized air stream was mixed with a carrier air stream that had a flow rate of 1000 mL/min. The diluted odor stream (1500 mL/min) was then split equally into two streams (750 mL/min each), one directed to the fly and the other directed to a photoionization detector (mini-PID, Aurora Scientific) to monitor odor delivery. PID signals were collected using the imaging software at 50 Hz. Air flow was turned on at least one minute prior to the start of imaging experiments. Odor delivery was controlled and synchronized with bitter delivery using MATLAB software.

### Immunohistochemistry

Brains were dissected in phosphate buffered saline (PBS) and fixed for 15-20 min in 4% paraformaldehyde in PBS. Brains were then washed multiple times with PBS containing 0.3% Triton X-100 (PBST), blocked with 5% normal goat serum diluted in PBS for 1 hr, incubated with primary antibodies at 4° for 2-3 days, washed in PBST, incubated with secondary antibodies at 4° overnight, then washed in PBST and PBS. Brains were mounted in Vectashield. Primary antibodies used were chicken anti-GFP (1:1000), rabbit anti-DsRed (1:500), and mouse nc82 (1:10). Secondary antibodies used were Alexa Fluor 488 goat anti-chicken (1:500), Alexa Fluor 568 goat anti-rabbit (1:500), and Alexa Fluor 633 goat anti-mouse (1:500). Images were acquired on a Zeiss LSM 710 system using a 25x or 40x objective. Confocal images were processed using Fiji/ImageJ.

## QUANTIFICATION AND STATISTICAL ANALYSIS

### Calcium imaging analysis

Calcium imaging data were analyzed using custom MATLAB code based on scripts from previous studies (Devineni et al., 2019; Hattori et al., 2017). In most experiments, images were registered within and across trials to correct for movement in the x-y plane using a sub-pixel registration algorithm. Regions of interest (ROIs) were drawn manually. Average pixel intensity within the ROI was calculated for each frame. The average signal for 20 frames preceding stimulus delivery was used as the baseline signal, and the ΔF/F (change in fluorescence divided by baseline) for each frame was then calculated. The peak ON response was quantified as the average ΔF/F value for the two highest consecutive frames during stimulus presentation. The peak OFF response was quantified in the same way for the 2.5 sec following stimulus offset, except that the ΔF/F value just prior to stimulus removal (average over last two frames) was then subtracted. The OFF response therefore represents how much the calcium activity increases upon stimulus offset. ON response adaptation was quantified as 1 – (average response on last two stimulus frames / peak ON response). Trials were excluded if the tastant drop failed to make proper contact with the leg or labellum based on video monitoring. In some experiments (noted in the figure legends) responses were not evoked on every trial and non-responsive trials were excluded from the averages.

For MBON imaging experiments the plasticity effect was calculated as follows: we separately averaged the CS+ and CS-responses across all blocks, then calculated the percent change from pre-pairing to post-pairing, and finally calculated the difference between these values for the CS+ and CS-. Because this method generates a single value for each experiment, we performed bootstrapping in MATLAB using 10000 iterations to obtain an estimate of the standard deviation.

### Decoding and statistical analysis

For decoding analyses, GCaMP traces were first normalized to the maximum response of each fly. ON and OFF responses were each quantified by multiple “features” representing the average response over different time windows. We compared decoding results using different numbers of features and found that just two features per response enabled decoding with similar accuracy as 4 or 8 features. These two features represented the “early” response (first 3 frames or ∼0.8 sec) and the “late” response (last two frames or ∼0.6 sec). Each feature was then scaled to the same range. Decoding analyses were performed using support vector classification with a linear kernel using the scikit-learn library in Python. Decoder performance was quantified using k-fold stratified cross-validation, which was iterated 1000 times to ensure data were randomly allocated into “folds” (i.e., training and testing sets), and the average accuracy across iterations was calculated. For decoding with shuffled labels, the same cross-validation analysis was performed for 100 random shuffles of the labels, except that allocation of data into each fold was only iterated 10 times per label shuffle (representing 1000 total iterations), and the average accuracy was calculated.

To determine whether the plasticity effect of each MBON imaging experiment was statistically significant (compared to zero), we performed permutation testing in MATLAB using 10000 permutations. All other statistical analyses were performed using GraphPad Prism, Version 4. Statistical tests and results are reported in the figure legends. To compare two groups, a t-test was used when sample sizes were similar and the non-parametric Mann-Whitney test was used when sample sizes were disparate. One- or two-way ANOVA was used to compare multiple groups with Tukey’s or Bonferroni’s post-tests, respectively. All graphs represent mean ± SEM unless otherwise specified. Sample sizes are listed in the figure legends and the number of trials and flies is noted.

## DATA AND CODE AVAILABILITY

MATLAB and Python scripts and data generated in this study are available upon request.

## SUPPLEMENTAL INFORMATION

Supplemental Information includes five figures and one table.

---

**Figure S1, related to Figure 1:**
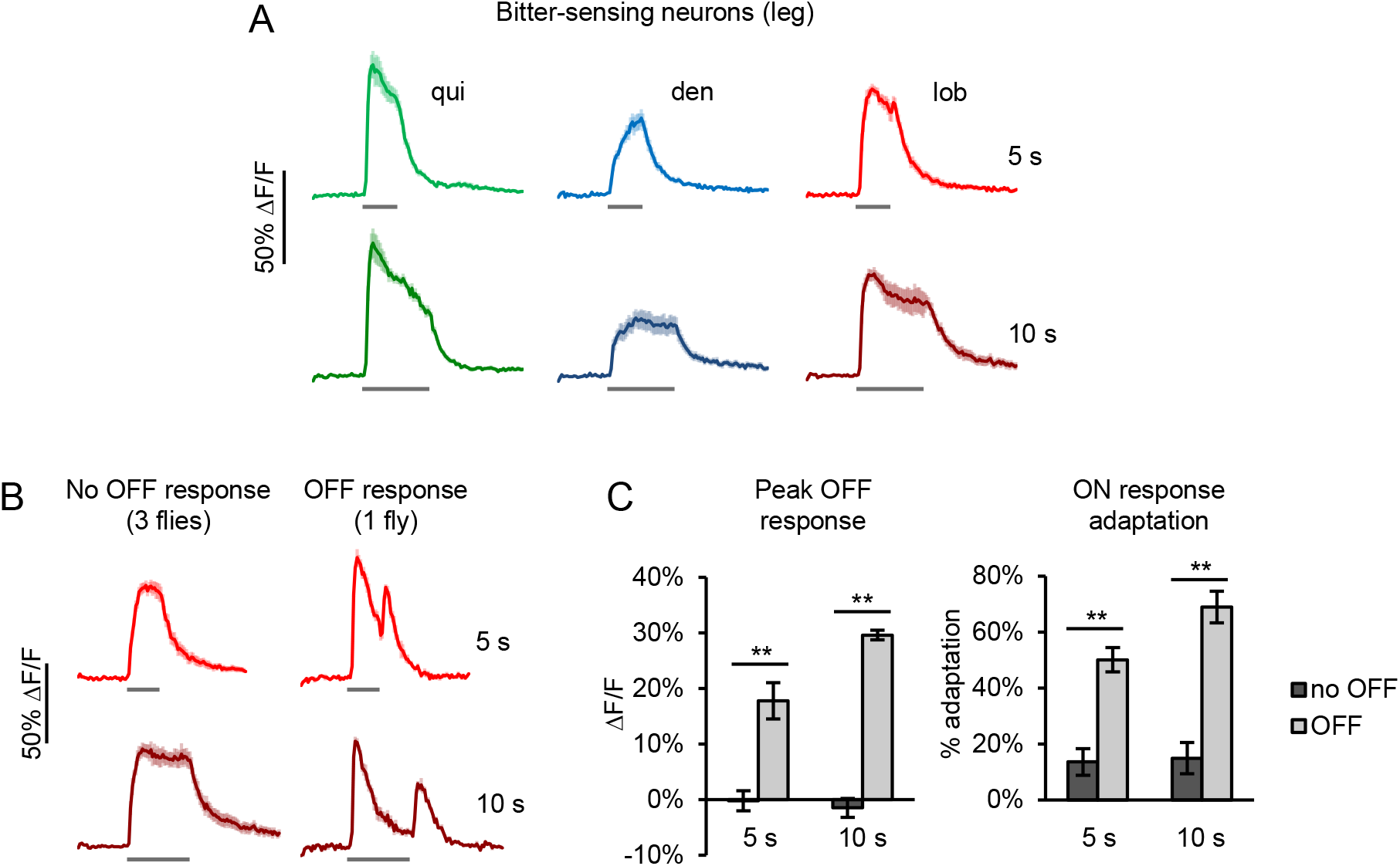
Additional imaging of bitter neurons in the leg. *Gr33a-Gal4* was used to label a larger number of bitter neurons in the foreleg than those labeled by *Gr98d-Gal4* in Figure 1 (Ling et al., 2014). (A) *Gr33a-Gal4-*expressing bitter neurons in the leg display ON responses but no OFF response to 5 sec bitter stimulation (n ≥ 12 trials, 4 flies). Non-responsive trials were excluded from these averages. (B-C) In 1 of 4 flies tested, tarsal bitter neurons showed a consistent OFF response to lobeline, but not to other compounds. Interestingly, the lobeline response in this fly also showed greater ON response adaptation. (B) Average lobeline response in this fly (right; n = 4 trials) compared to the 3 other flies (left; n = 9 trials). (C) Quantification of the peak OFF response and ON response adaptation in the fly showing a lobeline OFF response (“OFF”) and the three other flies (“no OFF”). Results for 5 sec or 10 sec lobeline stimulation are shown separately. **p<0.01, Mann-Whitney test.

**Figure S2, related to Figure 2:**
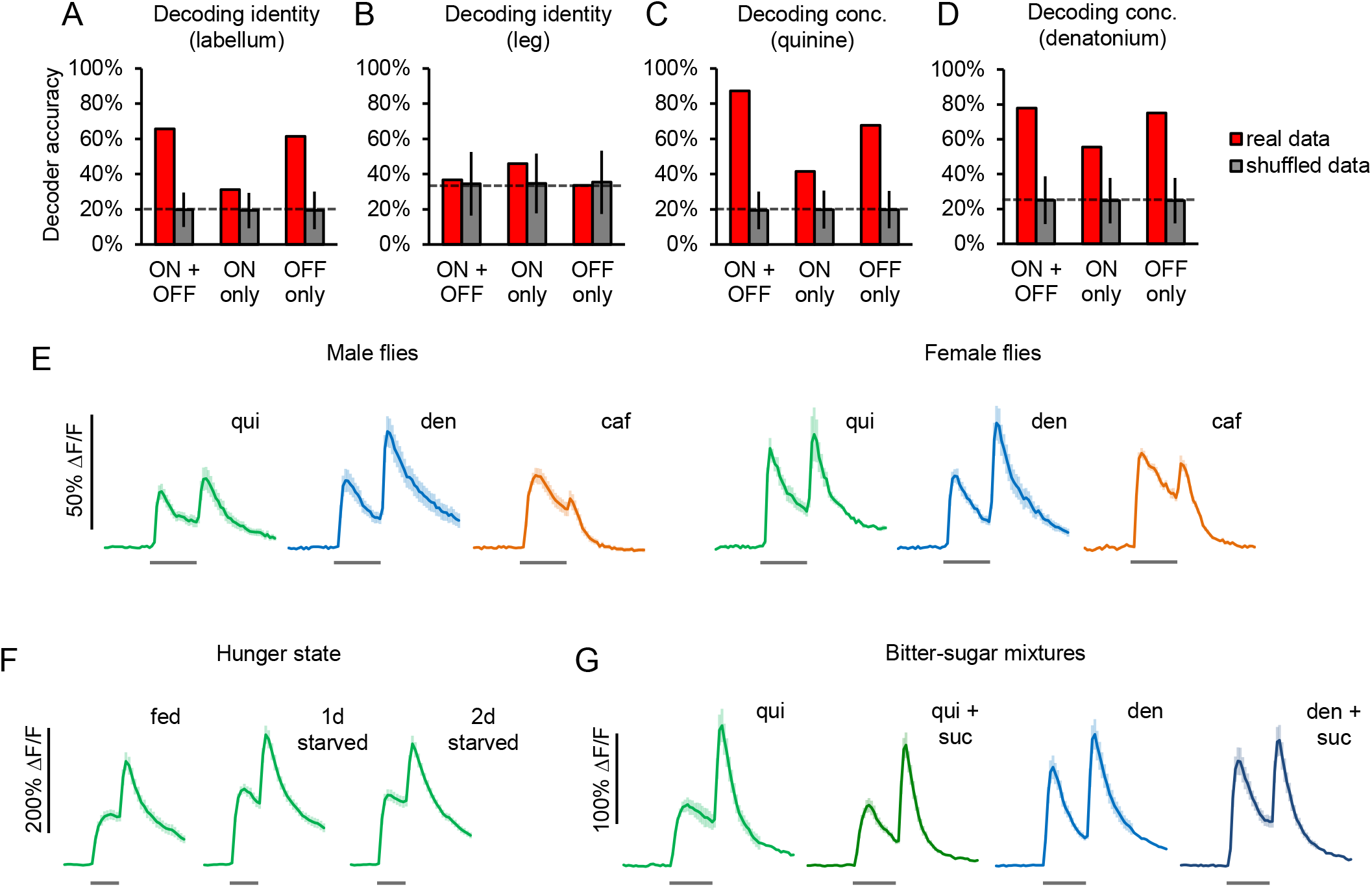
Bitter response dynamics encode stimulus identity and concentration but are not modulated by sex, hunger, or the presence of sugar. (A-D) Performance of decoders trained to classify stimulus identity (A-B) or concentration (C-D). For each dataset, separate decoders were trained using either the ON or OFF response only or both the ON and OFF responses. Decoding of data with shuffled labels was performed as a control. Dashed lines denote chance level based on number of groups (stimulus types or concentrations) and error bars represent 95% confidence intervals. (A) Decoder performance on classifying stimulus identity from labellar bitter responses (five compounds, data from Figure 2E-F). (B) Decoder performance on classifying stimulus identity from tarsal bitter responses (three compounds, data from Figure 1E). (C-D) Decoder performance on classifying the concentration of quinine (C; five concentrations) or denatonium (D; four concentrations) (data from Figure 2G-H). (E) *Gr33a-Gal4*-expressing labellar bitter neurons in males (n = 16 trials, 4 flies) and females (n = 8 trials, 2 flies) showed similar response dynamics. Bitter stimulation was for 5 sec. Note that all other experiments in this study used female flies. (F) *Gr66a-Gal4*-expressing labellar bitter neurons in fed and starved flies showed similar response dynamics to 3 sec quinine stimulation (n = 24 trials, 8 flies per group). (G) The addition of sucrose to quinine or denatonium did not affect the response dynamics of *Gr98d-Gal4*-expressing labellar bitter neurons (n = 15 trials, 5 flies).

**Figure S3, related to Figure 5:**
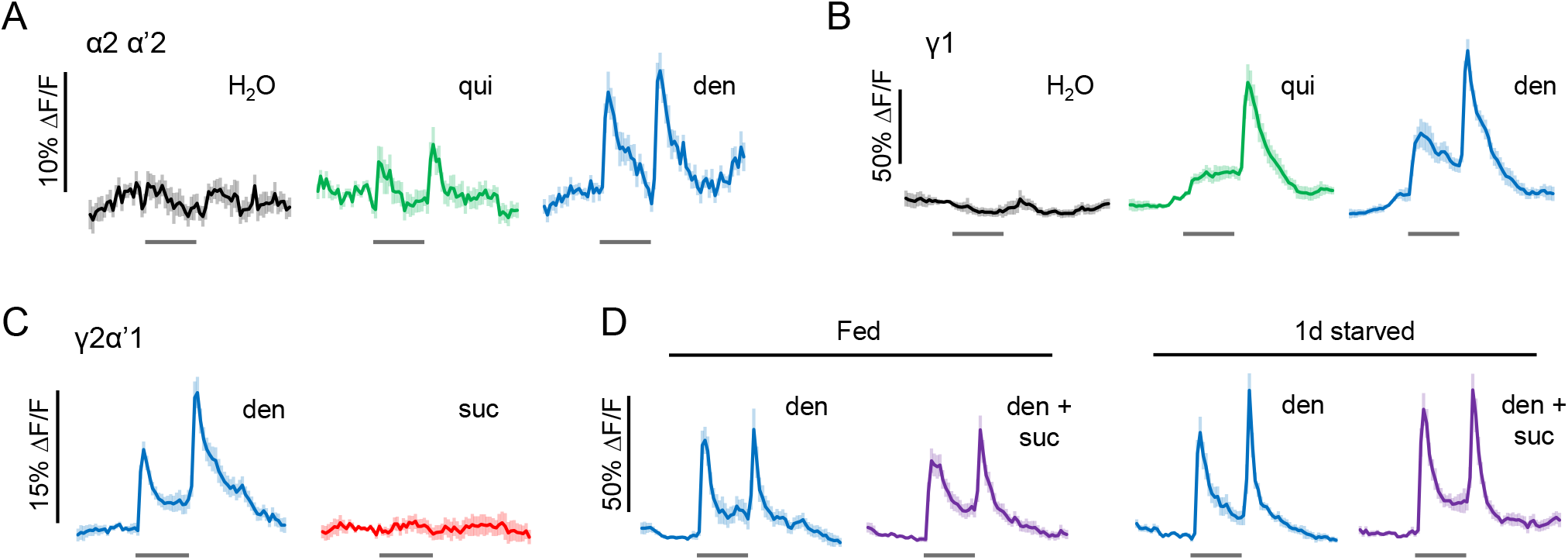
Further characterization of bitter responses in PPL1 DANs. (A-B) Bitter ON and OFF responses were observed in the α2 α’2 DAN (panel A; n = 11-16 trials, 3-4 flies) and the γ1 DAN (panel B; n = 16 trials, 4 flies). (C) The γ2α’1 DAN responded to bitter compounds such as denatonium but not to sucrose (n ≥ 38 trials, 7 cells, 5 flies). (D) The addition of sucrose to denatonium in fed or one-day starved flies did not affect the response dynamics of the γ2α’1 DAN (n ≥ 17 trials, 4 flies). The larger bitter responses in panel D compared to panel C are likely due to an improved imaging preparation that did not require removal of the antennae (see Methods). The α2 α’2 and γ2α’1 DANs were labeled by *TH-Gal4* (panels A, C-D). The γ1 DAN was labeled by the *MB320C* split *Gal4* line (panel B).

**Figure S4, related to Figure 6:**
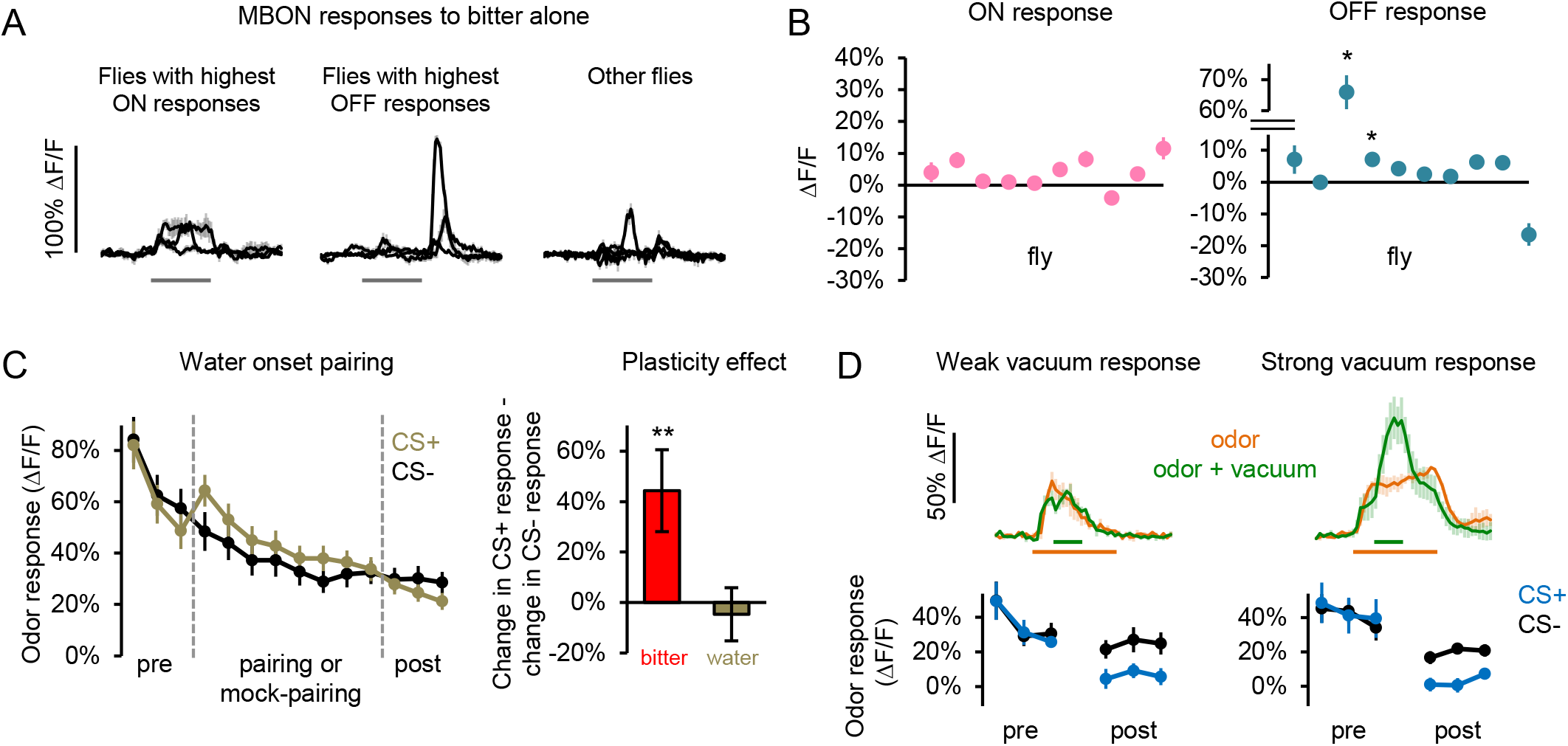
Controls for analyzing effects of odor-bitter pairing on MBON plasticity. (A-B) The γ2α’1 MBON was not reliably activated by bitter alone (5 sec denatonium). (A) Average GCaMP traces for individual flies (n = 4-5 trials). Separate graphs are plotted for the 3 flies with the highest ON response (left), 3 flies with the highest OFF response (middle, all different flies from first graph), and 4 additional flies. (B) Average ON and OFF responses for each fly (sorted arbitrarily). Responses were averaged over the first 1 sec of the ON or OFF period. *p<0.05, one-sample t-test compared to 0 followed by Sidak correction for multiple comparisons. (C) The onset of a water stimulus was paired with a CS+ odor using the same protocol as used for “bitter onset pairing” in Figure 6E-F (n = 7 flies, 20-21 blocks). Left: Mild facilitation of the CS+ response can be observed during the water-odor pairing, but this does not represent plasticity because CS+ and CS-responses did not differ post-pairing. Right: Water pairing did not induce significant plasticity. Plasticity for bitter onset pairing is shown for comparison, reproduced from Figure 6F. Error bars represent standard deviation obtained by bootstrapping, and p-values were obtained by permutation testing (**p<0.01). (D) During odor pairing with bitter offset, odor is also paired with the 1 sec vacuum pulse that removes bitter, which induced an MBON response in many flies. However, the vacuum pulse was also presented during mock-pairing trials in CS-blocks, so any effect of odor-vacuum pairing should affect CS+ and CS-responses equally. In addition, we separately analyzed flies whose MBONs showed weak (2 flies) versus strong (4 flies) vacuum responses, as determined during CS-mock-pairing trials. Top graphs: Average responses for the 3 pre-pairing trials (“odor”) and the first 3 mock-paired trials (“odor + vacuum”) for example flies in each group. Bottom graphs: Both groups of flies showed a similar magnitude of CS+ depression, suggesting that the vacuum response is not a major factor driving this plasticity (n = 2-4 flies, 6-12 blocks).

**Figure S5, related to Figure 7:**
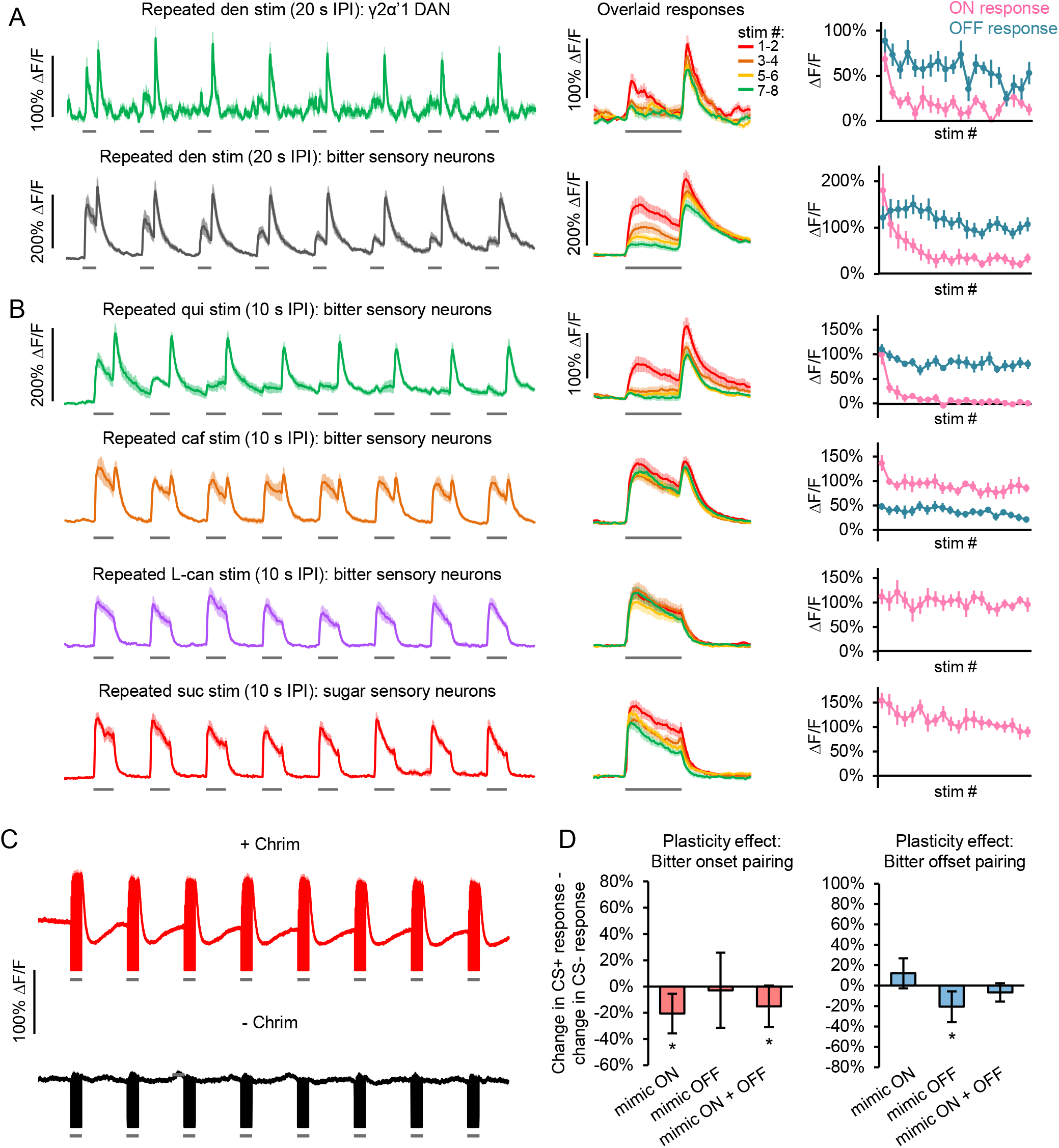
Further characterization of taste response habituation and controls for optogenetic pairing experiments. (A-B) Responses of the γ2α’1 DAN (labeled with *TH-Gal4*), bitter-sensing neurons (labeled with *Gr33a-Gal4*), or sugar-sensing neurons (labeled with *Gr64f-Gal4*) to repeated taste stimulation (5 sec each). Strong habituation of the ON response was observed for tastants that induce a strong OFF response (denatonium and quinine), but not other tastants. Left panels show average traces. Middle panels show responses to different stimulus repetitions overlaid, with baseline fluorescence calculated separately for each stimulus. Right panels show average ON and OFF responses to each stimulus. (A) Response to denatonium delivered with 20 sec IPI (n = 8-9 flies). (B) Responses of bitter neurons to repeated quinine, caffeine, or L-canavanine (top 3 rows) or response of sugar-sensing neurons to repeated sucrose (bottom row) delivered with 10 sec IPI (n = 6 flies). (C) Red light (grey bars) activated the γ2α’1 DAN in flies expressing Chrimson (top; n = 10 experiments, 5 flies), but not in control flies that did not express Chrimson (bottom; n = 8 experiments, 4 flies). 8 pulses of light were delivered in the same way as in the MBON imaging experiments (Figure 7G-H), with each pulse comprising 16 x 50 ms light. During light delivery the imaging shutter was closed, leading to a loss of GCaMP signal that is represented by the GCaMP traces falling well below baseline during each light pulse. (D) MBON plasticity effects for control flies that did not express Chrimson. Odor-light pairing experiments were performed in the same way as shown in Figure 7G.

**KEY RESOURCES TABLE**

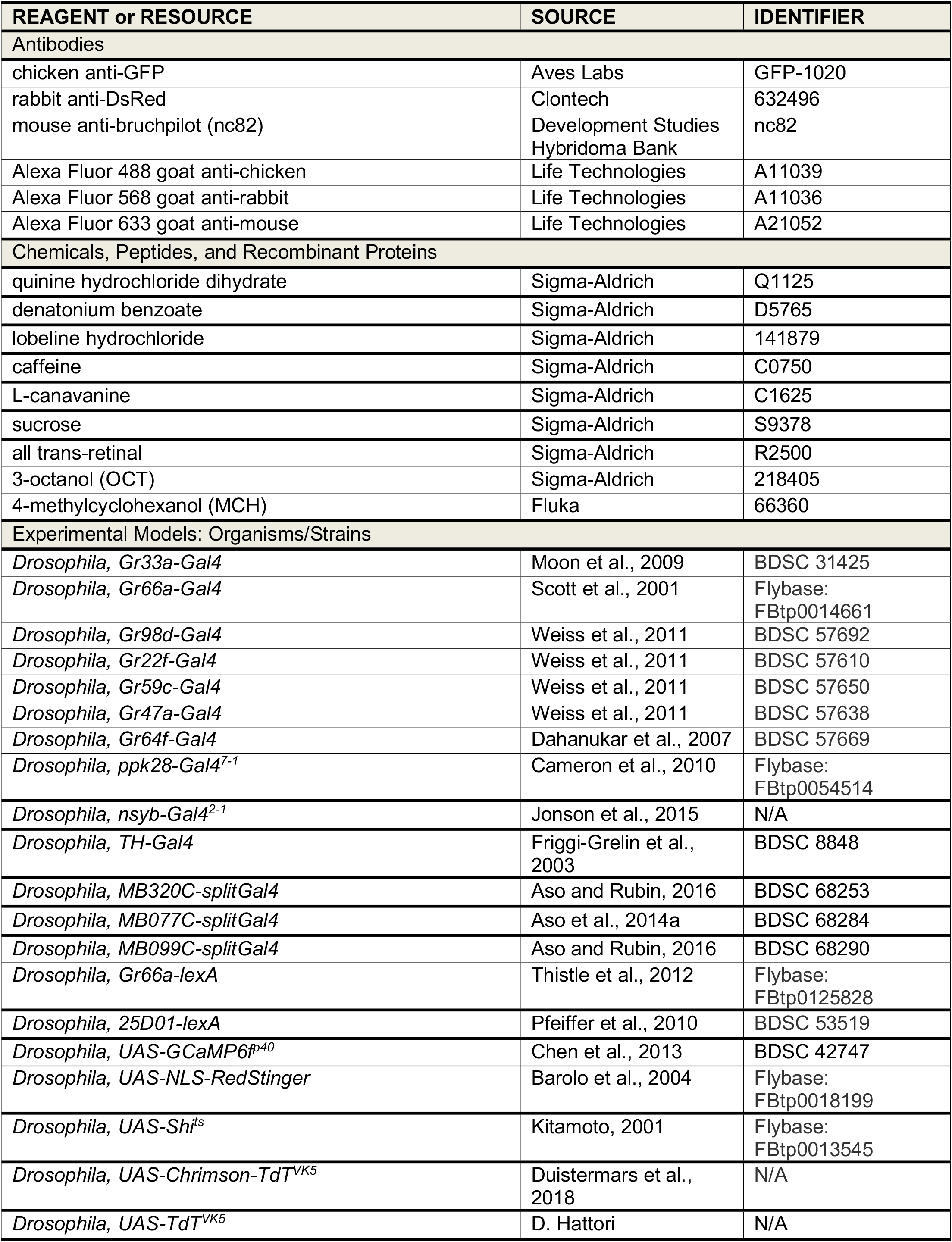
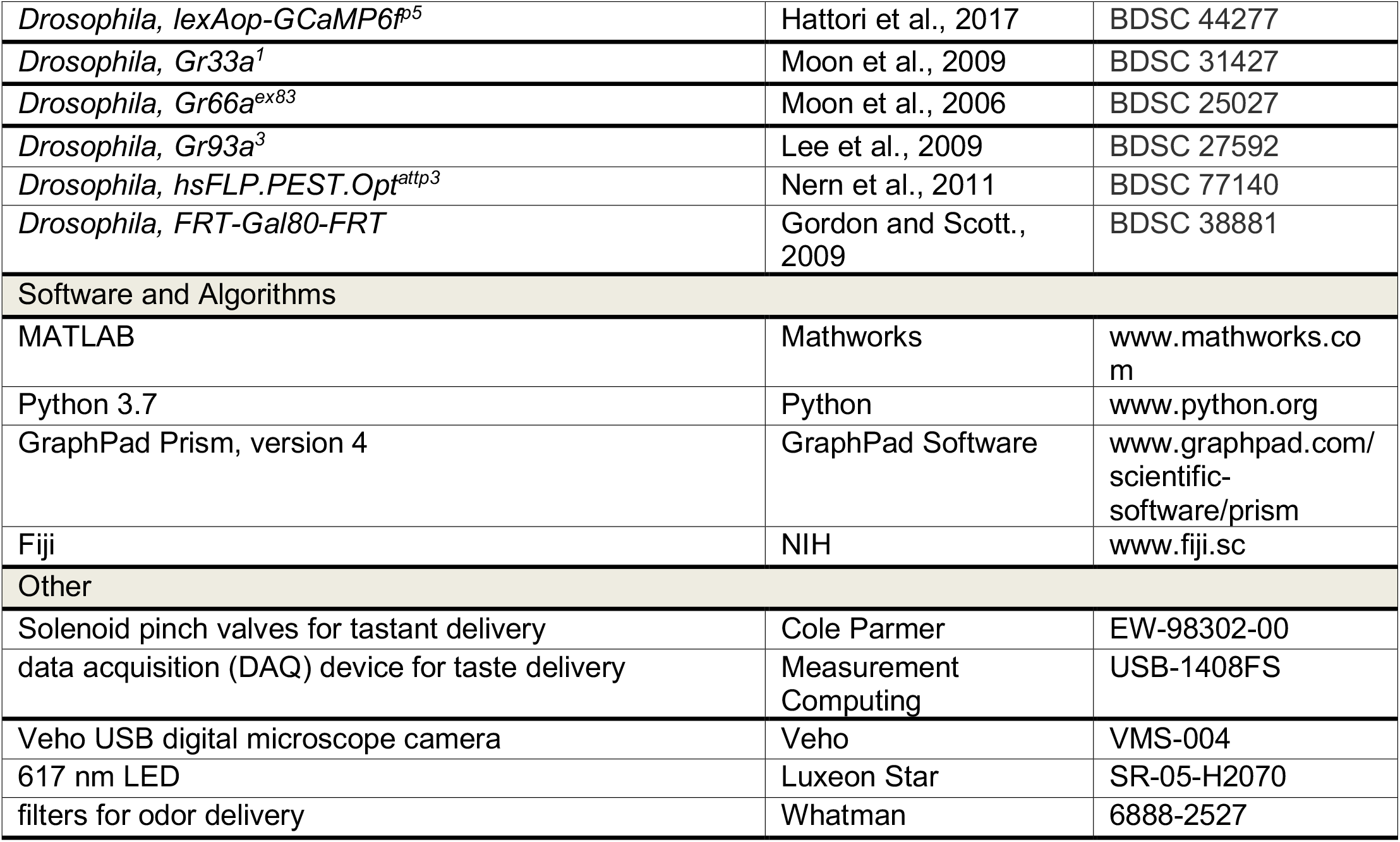

